# LassoESM: A tailored language model for enhanced lasso peptide property prediction

**DOI:** 10.1101/2024.10.25.620295

**Authors:** Xuenan Mi, Susanna E. Barrett, Douglas A. Mitchell, Diwakar Shukla

## Abstract

Ribosomally synthesized and post-translationally modified peptides (RiPPs) comprise a structurally and functionally diverse group of natural products. Lasso peptides represent one of about 50 known molecular classes of RiPPs, which display a characteristic [1] rotaxane conformation formed by a lasso cyclase. This unique, threaded conformation endows lasso peptides with diverse biological activities and remarkable thermal and proteolytic stability. The prediction of lasso peptide properties, such as substrate compatibility with a particular lasso cyclase or desired biological activity, remains challenging due to limited experimental data and the intricate nature of the substrate fitness landscapes. Protein language models (PLMs) have demonstrated impressive performance in predicting protein structure and function. However, general-purpose PLMs perform poorly in lasso peptide-related predictive tasks. Therefore, there is a need to provide effective representations for lasso peptides to enable enhanced property prediction. In this study, we developed a lasso peptide-specific language model (LassoESM) by leveraging advances in pre-trained PLMs to aid the prediction of lasso peptide related properties and experimentally validate the model predictions. We demonstrate that LassoESM embeddings can accurately predict substrate compatibility for a lasso cyclase of interest when experimental data for model training was scarce. Using a deep learning framework incorporating cross-attention between lasso cyclase and substrate peptide embeddings, we identify non-cognate pairs of lasso cyclases and substrate peptides with predicted compatibility. We further show that LassoESM embeddings improve the prediction of RNA polymerase inhibitory activity, which represents a biological activity of several known lasso peptides. We anticipate that LassoESM and future iterations thereof will be instrumental for the rational design of lasso peptides with desired properties.

## Introduction

Ribosomally synthesized and post-translationally modified peptides (RiPPs) are structurally diverse natural products. RiPP precursor peptides are genetically encoded and many biosynthetic enzymes are tolerant to alternative substrates, making RiPPs attractive engineering targets for a variety of medical and other applications.^1^ Lasso peptides are a particularly promising scaffold for RiPP engineering due to their modifiable and stabilized lariat knot-like structures.^1,2^ During lasso peptide biosynthesis, the precursor peptide is bound by the RiPP recognition element, and a leader peptidase releases the core region of the substrate that undergoes conversion into the mature product. An ATP-dependent lasso cyclase then folds the core peptide into its characteristic threaded shape, which is kinetically trapped by macrolactam formation (**Figure 1A**).^3,4^ The N-terminal amine (core position 1) and the side chain of Asp or Glu, typically located at core position 7, 8, or 9, form the macrolactam with large residues in the tail acting as steric locks to maintain threadedness.

**Figure 1.**
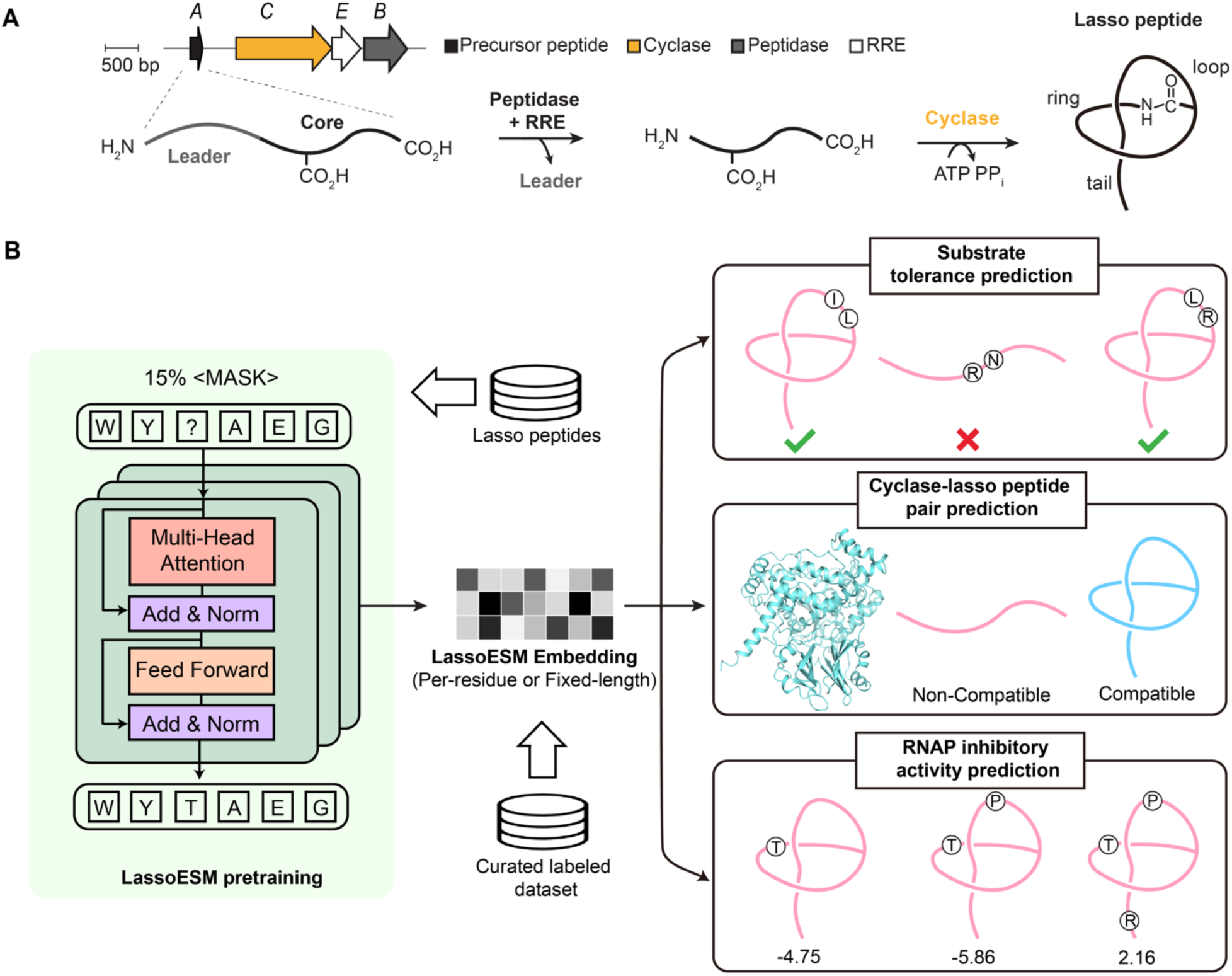
Development of a lasso peptide-specific language model, LassoESM. **(A)** Lasso peptide biosynthesis requires a leader peptidase, RiPP recognition element (RRE), and lasso cyclase to tie a linear core peptide into the lariat-like knot. **(B)** LassoESM was built upon the ESM-2 architecture and further pre-trained on lasso peptides using a domain-adaptive approach with masked language modeling. The resulting LassoESM embeddings were utilized for three downstream tasks: predicting lasso cyclase substrate tolerance, identifying substrate compatibility between non-cognate pairs of lasso cyclases and substrate peptides, and predicting RNAP inhibitory activity.

Several lasso cyclases have shown high substrate tolerances, such as the fusilassin cyclase (FusC: WP_104612995.1), which was statistically shown to cyclize millions of variants with multi-site changes to the ring,^5^ and the microcin J25 cyclase (McjC: WP_256498469.1), which can accommodate many diverse single-site and multi-site changes.^6,7^ Additionally, lasso peptides exhibit antibacterial, antiviral, and anticancer activities, making them a compelling scaffold for engineering macrocyclic peptides for drug discovery purposes.^4,8^ Despite their potential, chemical synthesis of lasso peptides remains highly challenging, with only one successful report that has yet to be replicated.^9^ Therefore, designing novel lasso peptides with desired bioactivities will require sequences that are compatible with available lasso cyclases. However, profiling the substrate tolerance of lasso peptide biosynthetic enzymes remains challenging, hampering the generation of customized lasso peptides for biomedical and other industrial applications.

Different *in silico* approaches can be utilized to discover and design novel lasso peptides. For instance, RODEO (https://webtool.ripp.rodeo/) predicts the native precursor peptides that are proximally encoded to the lasso peptide biosynthetic enzymes using linear combination scoring and a supervised machine learning approach.^10^ RODEO matches substrates with enzymes but cannot predict whether a biosynthetic enzyme would tolerate an alternative peptide substrate. Additionally, RODEO only predicts precursor peptides and their associated biosynthetic enzymes within a small local genomic context, which precludes the possibility of identifying potential hybrid peptide-enzyme pairs. Molecular dynamics simulation is a powerful computational approach for studying how lasso peptides or other RiPPs fold and how they interact with their biosynthetic enzymes during folding.^11,12,13^ However, molecular dynamics simulations are computationally expensive, limiting their applicability for the broadscale customization of lasso peptides with desired properties.

Deep learning is a type of artificial intelligence that can learn complex patterns and features from large datasets. It has emerged as a useful tool for predictive tasks related to RiPP-modifying enzymes, including predicting RiPP enzyme substrate tolerance and bioactivity retention. For example, a convolutional neural network classifier was trained to predict substrate preferences for the lactazole pathway (an example of a thiopeptide).^14,15^ These methods enabled the design of a diverse and highly modified variant library for selection against a target. As a second example, DeepLasso predicts whether a variant of the lasso peptide ubonodin will retain RNA polymerase (RNAP) inhibitory activity.^16^ The model demonstrates the utility of artificial intelligence beyond substrate compatibility prediction for lasso peptide biosynthesis. However, these methods required uncharacteristically large datasets for model training. In most cases, the experimental data that defines lasso peptide substrate tolerance and bioactivity is severely limited, necessitating methods that perform well in data scare scenarios.

Protein language models (PLMs) are a powerful framework for improving the performance of downstream tasks in situations with limited labeled data.^17,18,19,20,21,22^ Most PLMs are trained using masked language modeling, in which the model is trained to predict the identity of masked amino acids given the local amino acid sequence. PLMs have been used to improve predictions for protein structure,^22^ protein properties,^23,24,25^, and mutational effects.^17,24,26,27^ Effective PLMs require massive training datasets and large model size, which enable learning of general patterns present in natural proteins. However, PLMs are predominantly trained on protein sequences rather than peptides, and their ability to represent peptides is understudied. Since peptides differ from proteins in physical characteristics, such as shorter length, simpler three-dimensional structures, and conformational dynamics, we hypothesized that PLMs may have limited abilities to accurately represent peptide sequences, particularly lasso peptides, which exhibit a unique knot-like shape. As a result, the effectiveness of PLMs in performing predictive tasks related to lasso peptides appears to be a logical mismatch. Previous studies have demonstrated that domain-adaptive pretraining can further enhance the performance of large language models within a specific domain by learning the knowledge of the given domain.^28,29,30^ For example, pretraining on DNA-binding protein sequences has improved model performance in four DNA-binding, protein-related downstream tasks.^29^ Similarly, pretraining on a dataset focused on the substrate preference of one enzyme can enable accurate prediction of which substrates will be compatible with another enzyme in the same biosynthetic pathway.^30^ Due to the diversity of RiPP classes and their unique sequence attributes, it is possible that class-specific RiPP language models could further improve downstream predictive tasks.

In this work, we used a domain adaption method to further pre-train ESM-2 (Evolutionary Scale Modeling) on lasso peptide datasets, resulting in a lasso peptide-tailored language model named LassoESM. Compared to the more generic ESM-2, LassoESM showed superior performance in predicting the substrate tolerance of a known lasso cyclase, even with small quantities of training data. A Human-in-the-Loop approach, where a human expert iteratively validates new, unseen sequences and incorporates them into the training data, was used to further verify and improve the classification model.^31^ To predict compatible lasso cyclase-substrate peptide pairs, LassoESM embeddings were combined with a cross-attention layer to effectively model the interactions between the lasso cyclase and peptide, leading to better model performance than when only the generic PLM was used. Finally, LassoESM embeddings were shown to predict lasso peptide RNAP inhibition activity more accurately than a generic PLM and DeepLasso.^16^ With expanded lasso peptide property datasets, LassoESM could facilitate their biomedical application by guiding lasso cyclase selection for rational engineering campaigns.

## Methods

### Dataset construction for LassoESM

A bioinformatics tool, RODEO, was developed to automate the identification of RiPP biosynthetic gene clusters (BGCs), including lasso peptide BGCs, from genomes available in GenBank.^10^ RODEO utilizes profile Hidden Markov Models and a support vector machine classifier to locate RiPP BGCs and predict the most likely precursor peptide, respectively. A previous study identified 7,701 non-redundant, high-scoring lasso precursor peptides using RODEO.^5^ After removing identical entries (if two distinct organisms encode the same core sequence, only one was retained), 4,485 unique core sequences were retained for pre-training LassoESM.

### Model architecture and Pre-training

ESM-2 is a PLM trained on ∼65 million non-redundant protein sequences using a masked language modeling approach.^22^ Masked language modeling has been shown as a powerful pretraining technique for language models. In our work, we adapted the ESM-2 (with 33 layers and 650 million parameters) architecture as the starting point for pretraining on lasso peptide datasets. During pre-training, 15% of the amino acids in the sequences were randomly masked. The model was trained to predict the identity of masked amino acids based on the surrounding sequence context, as shown in Equation (1):

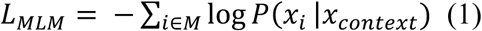

where M represents a set of indices of the randomly masked amino acids. The model was optimized to minimize the negative likelihood of true amino acid *x_i_* given the surrounding context *x_context_* in the lasso peptide sequences. By updating the full 650 million parameters of the ESM-2 model during pretraining, we enabled the model to learn specialized sequence patterns unique to lasso peptides, named LassoESM.

LassoESM was trained for 20 epochs with a learning rate of 5 ×10^−5^ and a batch size of 8 on a single NVIDIA GeForce RTX 3090 GPU with 12 GB memory. To reduce the memory required, GaLore (Gradient Low-Rank Projection) was integrated into the training process.^32^ GaLore is a training strategy that allows full-parameter learning but can significantly reduce the memory requirements traditionally associated with optimizer states and gradients during the training process. During training, the loss to be minimized is the mean cross-entropy loss between the predicted and the ground truth masked amino acid. The epoch with the least validation loss was saved as the final model. LassoESM is publicly available on Hugging Face (https://huggingface.co/ShuklaGroupIllinois/LassoESM).

### Fusilassin labeled dataset construction

The fusilassin substrate tolerance dataset contained substrate and non-substrate sequences confirmed via cell-free biosynthesis (CFB) and matrix-assisted laser desorption/ionization time-of-flight mass spectrometry (MALDI-TOF-MS) from the following previously published libraries: 5 × NNK ring library,^5^ 5 × NNK loop library,^5,13^ and three 2 × NNK loop panels (*note*: NNK encodes for all 20 common proteinogenic amino acids and a stop codon).^13^ The dataset also contained a variety of published and unpublished variants of fusilassin. All sequences were verified as substrates or non-substrates using CFB with purified enzymes: 10 µM each of maltose-binding protein (MBP)-FusB, MBP-FusE, and MBP-FusC. The reactions were analyzed for lasso cyclization with MALDI-TOF-MS.

### Training of downstream models

#### Lasso peptide substrate tolerance prediction

Three language models were used in this study: VanillaESM (650 million parameters ESM-2),^22^ PeptideESM^30^, and LassoESM. The last layer of these language models was utilized to generate information-rich representations, also known as embeddings. For each amino acid in each labeled lasso peptide sequence, the last layer of the language model generated a 1,280-dimensional vector. The final embeddings for each peptide sequence were obtained by calculating the average of all the amino acid vectors, which were then fed into downstream models.

A total of 1,121 fusilassin variant sequences were collected and experimentally labeled as substrates (656 sequences) or non-substrates (465 sequences). Embeddings were generated using one-hot encoding, VanillaESM, PeptideESM, and LassoESM to represent these sequences. The embeddings were then used to train various downstream classification models, including Random Forest (RF), Adaptive Boosting (AdaBoost), Support Vector Machine (SVM), and Multi-Layer Perceptron (MLP). Stratified 10-fold cross-validation was employed to optimize the hyperparameters of supervised classification models. The optimized hyperparameters for all downstream models are in Table S1. The model performance was evaluated using AUROC and balanced accuracy, with both metrics calculated from 5 repeats of 10-fold cross-validation. All downstream classification models were implemented using Scikit-learn.^33^

The same approach was applied to microcin J25 (MccJ25) variant sequences to predict the substrate tolerance for the microcin J25 cyclase. In this case, 552 MccJ25 variants were collected from the literature, with 413 sequences labeled as substrates and 139 as non-substrates.^6,7,34,35,36,37,38^ The optimized hyperparameters for all downstream models are in Table S2.

#### Prediction of lasso cyclase-substrate peptide pairs

A total of 6,599 unique lasso cyclase-substrate peptide pairs were identified using RODEO.^10^ To generate synthetic, non-natural, cyclase-peptide pairs as negative samples, we first generated all possible non-natural pairs for each cyclase. From these, we retained only those pairs in which the acceptor residue of the peptide differed from the original acceptor residue, as the acceptor residue is the most restrictive and is exclusively either Asp or Glu (only two examples are known where a lasso cyclase tolerates Asp and Glu^39,40^). Finally, one lasso peptide was randomly selected from this pool for each cyclase, generating 6,599 synthetic non-natural cyclase-peptide pairs as negative samples.

LassoESM was used to generate embeddings for lasso peptides, while VanillaESM was employed to generate embeddings for lasso cyclases. For each amino acid in the lasso peptide sequence, the last layer of LassoESM produced a 1,280-dimensional vector, resulting in a matrix of dimensions *d_1280_* ∗ *m*, where *m* is the length of the lasso peptide. For each amino acid in the lasso cyclase sequence, the last layer of VanillaESM generated a 1,280-dimensional vector, resulting in a matrix with *d_1280_* ∗ *n*, where *n* is the length of the lasso cyclase. Padding was added at the end of the substrate peptide and lasso cyclase sequences to ensure that all substrate peptide embeddings had the same dimensions and that all lasso cyclase embeddings had the same dimensions.

These two matrices were then supplied into a cross-attention layer, where the lasso peptide embedding matrix served as the query and the lasso cyclase embedding matrix as the key and value. An attention mask was applied to every padding position, as well as the BOS (Beginning of Sentence) and EOS (End of Sentence) tokens, to ensure the model ignored these elements. By performing a dot product between the query and key, an attention matrix was obtained, which was used to reweight the cyclase embedding matrix. The reweighted cyclase matrix was then concatenated with the lasso peptide matrix and fed into an MLP model. The MLP model consisted of two hidden layers with 512 and 64 neurons. The full model, including the cross-attention layer, was trained using the Adam optimizer, with a batch size of 32 and a learning rate of 0.001. The training was conducted for 25 epochs using PyTorch.^41^

#### RNA polymerase inhibition prediction

A total of 11,363 published ubonodin variant sequences with a measurable enrichment value were collected.^16^ After removing sequences containing stop codons, 8,885 ubonodin variants remained in the dataset. The last layers of VanillaESM, PeptideESM, and LassoESM were used to extract embeddings for each amino acid in the ubonodin sequences. The final embedding for each sequence was obtained by averaging the embeddings of all amino acids in the sequence. These averaged embeddings were then utilized to train an MLP regressor. The MLP model consisted of two hidden layers with 256 and 32 neurons. MLP was trained using the Adam optimizer, with a batch size of 16 and a learning rate of 0.001. The model was trained for 100 epochs using PyTorch, ^41^, and early stopping was applied to monitor the validation loss.

A similar approach was applied to another lasso peptide, klebsidin. A total of 340 klebsidin variant sequences were obtained from a previous publication.^42^ The embeddings of these sequences were supplied to various downstream regression models, including RF, AdaBoost, SVM, and MLP. Among these, the AdaBoost model achieved the best performance on the klebsidin dataset, as determined by the average R² score from 10-fold cross-validation. The optimal hyperparameters for AdaBoost were n_estimators = 200 and learning_rate = 1.

### Experimental materials and methods

Synthetic DNA primers and ultramers were ordered from Integrated DNA technologies. New England Biolabs supplied the Q5 DNA polymerase and the Gibson assembly mix. Chemical reagents were from Sigma-Aldrich unless otherwise noted. MALDI-TOF-MS data was acquired at the University of Illinois Urbana-Champaign in the School of Chemical Sciences on a Bruker UltraXtreme instrument in reflector positive mode. The MALDI-TOF-MS data was analyzed using Bruker FlexAnalysis software. Sequencing was performed at the Core DNA Sequencing Facility at the University of Illinois at Urbana-Champaign.

### Molecular biology techniques

The fusilassin biosynthetic genes were previously cloned into a modified pET28 vector that provides an N-terminal MBP tag for solubility and stability.^13,43^ This vector was used for the expression of the fusilassin leader peptidase (FusB, WP_011291590.1), RiPP Recognition Element (RRE, FusE, WP_011291591.1), and an *E. coli* codon-optimized cyclase (FusC_op_, WP_104612995.1). A previously reported pET28-*mbp-fusA* (precursor peptide) construct was used as a template to prepare all the fusilassin variants needed for this study.^43^

### Protein purification

Previously described methods were used to express and purify MBP-tagged FusB, FusE, and FusC_op_.^13,43^ Protein concentrations were determined with a Nanodrop OneC instrument measuring the absorbance at 280 nm and by Bradford assay. Extinction coefficients were calculated using the Expasy ProtParam tool.^44^ Protein purity was determined by visual inspection of a Coomassie-stained gel after separation via SDS-PAGE (Figure S12).

### Overlap extension PCR to construct fusilassin library members tested in CFB

DNA templates encoding the FusA variants were prepared by overlap extension PCR.^45^ Primers CFB_F and CFB_R were used to amplify the completed linear DNA product (Table S12). DNA concentration was determined by absorbance at 260 nm using a Nanodrop OneC instrument.

### Generation of linear DNA for chimeric lasso peptide sequences tested in CFB

Ultramer oligos encoding the fusilassin leader peptide and the desired core peptide were amplified by PCR and cloned into the pET28-*mbp* vector using Gibson assembly (Table S12). After sequence confirmation by Sanger sequencing, the linear DNA encoding the MBP-tagged chimeric peptide was amplified using primers CFB_F and CFB_R. The DNA concentration was determined on a Nanodrop OneC instrument at an absorbance of 260 nm.

### Cell-free biosynthesis reagent preparation

*E. coli* cellular lysates, energy mix, and GamS were prepared according to Kretsch et al.^46^

### Cell-free biosynthesis reactions

Reaction conditions from Si et al.^5^ were used with the following changes: 200 ng of linear DNA template per 20 µL reaction and a final concentration of 10 µM each of recombinantly expressed and purified MBP-FusB, MBP-FusE, and MBP-FusC_op_.

## Results and Discussion

### LassoESM improves lasso peptide substrate tolerance prediction

Representative PLMs like ESM^22^ and ProtBERT,^18^ have been trained on general protein sequences to learn sequence patterns across natural proteins in a self-supervised manner. Compared to general proteins, lasso peptides are much shorter (15-20 amino acids) and have a unique lariat knot-like structure.^1,4^ The development of a lasso peptide-specific language model has the potential to enhance the lasso peptide representation capabilities and improve the performance of downstream predictive tasks. To address this, we adapted the ESM-2 architecture^22^, which contains 650 million parameters, as a starting point and further trained it on a dataset of 4,485 unique, high-scoring lasso core peptides identified by RODEO. Using the masked language modeling approach, we developed LassoESM, a specialized language model tailored to lasso peptides. We then utilized LassoESM embeddings in various downstream lasso peptide-related predictive tasks, including predicting lasso cyclase substrate tolerance, identifying non-natural cyclase-substrate peptide pairs, and predicting RNA polymerase inhibitory activity of lasso peptides.

Previous studies have shown that FusC exhibits a range of substrate tolerability, from promiscuous to highly intolerant, depending on the specific core position being varied.^5,13,43^ However, the rules that govern substrate compatibility are poorly understood, making substrate tolerance prediction challenging. A total of 1,121 fusilassin (FusA) variants were confirmed as substrate or non-substrate sequences via CFB (**Figure 2A**). This dataset was selected to evaluate if LassoESM could improve the substrate tolerance prediction for FusC. Embeddings for all 1,121 sequences generated by LassoESM were fed into various downstream classification models, including RF,^47^ AdaBoost,^48^ SVM,^49^ and MLP.^50^ Hyperparameter optimization was conducted using 10-fold cross-validation (**Table S1**), and the performance of the downstream classifiers was evaluated by AUROC score and balanced accuracy. Previously, we developed PeptideESM, a peptide-specific language model trained on 1.5 million unique peptide sequences.^30^ We evaluated PeptideESM embeddings for lasso peptide substrate tolerance prediction. Among these models, SVM demonstrated the highest performance (**Figure 2B, Figure S1**). Across the various downstream classification models, PeptideESM and LassoESM provided superior embeddings for FusC substrate specificity prediction compared to that from the general protein language model (VanillaESM) and the one-hot baseline representation method.

**Figure 2.**
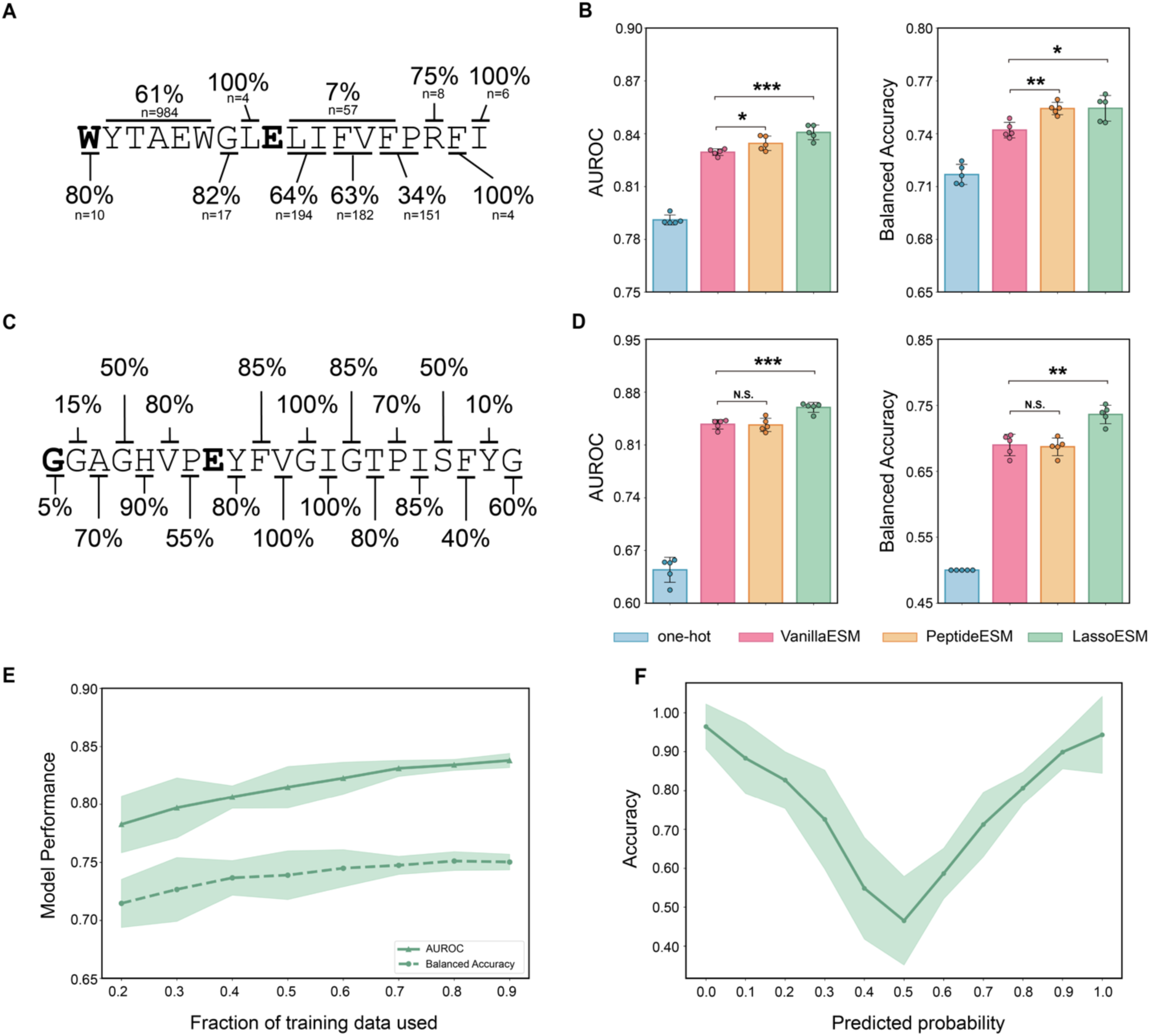
Comparison of lasso peptide substrate tolerance prediction using different embeddings. **(A)** Core peptide sequence of fusilassin overlaid with substrate tolerance information. The n value is the number of sequences tested at the indicated position(s) by CFB, with the percentage of n sequences tolerated by the biosynthetic enzymes given. Bold indicates the macrolactam-forming residues. **(B)** AUROC score and balanced accuracy of the SVM model for the fusilassin variant dataset trained on embeddings from one-hot encoding (blue), VanillaESM (pink), PeptideESM (orange) and LassoESM (green). Error bars: standard deviation calculated over 5 repeats of 10-fold cross-validation. *p*-value was calculated using a two-sided t-test. \**p* < 0.05, ** *p* < 0.01, *** *p* < 0.001. **(C)** Core peptide sequence of MccJ25 overlaid with substrate tolerance information. Each position was site-saturated with all other proteinogenic amino acids (n = 20) prior to heterologous expression.^6^ The percentage of sequences that were cyclized and exported is given. Bold indicates the macrolactam-forming residues. **(D)** Same as (**B**), except for MccJ25. **(E)** AUROC score and balanced accuracy of the SVM model trained on a fraction of the fusilassin variant training data using LassoESM embeddings and evaluated on the remaining data. Error bars: standard deviation across 10 random seeds, with a fraction of the dataset randomly selected for training in each seed. **(F)** Model accuracy at different predicted probabilities of the SVM model trained on the fusilassin variant dataset using LassoESM embeddings. Error bars: standard deviation calculated over 10 random seeds. For each random seed, 80% of the fusilassin variant dataset was used for training and the remaining 20% for testing.

Microcin J25 (MccJ25) is a distinct lasso peptide whose biosynthetic enzymes are reported to be highly substrate-tolerant. A total of 552 MccJ25 variants have been experimentally evaluated as substrates or non-substrates in the literature (**Figure 2C**).^6,7,34,35,36,37,38^ Analogous to fusilassin, LassoESM significantly outperformed VanillaESM and one-hot encoding to predict the substrate tolerance of McjC (**Figure 2D**) and SVM displayed the highest performance (**Figure 2D, Figure S2**). These results show that LassoESM provides effective representations that enhance the accuracy of substrate tolerance predictions for two disparate lasso peptide cyclase, FusC and McjC, which have a sequence identity of only 22%.

An advantage of language models for performing downstream classification tasks is their ability to improve accuracy in data-scarce settings.^51^ Researchers often lack access to large, labeled datasets for various biochemical tasks. This is especially true for RiPP biosynthetic enzyme compatibility prediction, given the overwhelming size of the sequence space. Thus, we evaluated the accuracy of the best downstream classification model, SVM, trained on datasets of differing sizes using LassoESM embeddings. For each training dataset size, we conducted 10 random selections of samples at different random seeds and evaluated the AUROC score and balanced accuracy on the remaining data samples. Even with only 20% of the training dataset (224 samples), the average AUROC score was 0.78, and the average balanced accuracy was 0.71 (**Figure 2E**). The results indicated that the classification model performs well even with a small dataset when using tailored language model embeddings. However, expanding the dataset with more diverse samples enhances model performance.

In classification tasks, the predicted probability is often interpreted as a measure of model uncertainty. However, the output from SVM is the distance from the decision hyperplane, not a probability. To address this, Platt scaling^52^ is commonly employed, where a logistic regression model is applied to the SVM decision values, effectively transforming distances into probability estimates ranging from 0 to 1. These probabilities represent the likelihood of a sequence being a substrate for the lasso cyclase. To assess whether these predicted probabilities effectively captured the uncertainty of the downstream classification model trained on the fusilassin dataset, we analyzed the distribution of the model accuracy across different probability ranges. Specifically, we plotted the accuracy distributions for the SVM models trained on LassoESM embeddings for fusilassin variant sequences (**Figure 2F**). The results indicated that the model demonstrated high accuracy when the predicted probabilities were near 0 or 1. Conversely, predictions with probabilities between 0.4 and 0.6 exhibited a higher proportion of false classifications. This result suggested that the predicted probability was faithfully representing model uncertainty.

### Human-in-the-Loop feedback improves fusilassin substrate specificity prediction

Next, we assessed the accuracy of LassoEM in predicting the substrate tolerance of sequences not included in the training set. A human-in-the-loop strategy was employed to further improve model performance.^31^ The workflow begins by classifying new sequences using the optimized SVM, trained on LassoESM embeddings from an initial dataset of 1,121 fusilassin variant sequences. This process is followed by human expert validation to enhance predictions (**Figure 3A**) iteratively.

**Figure 3.**
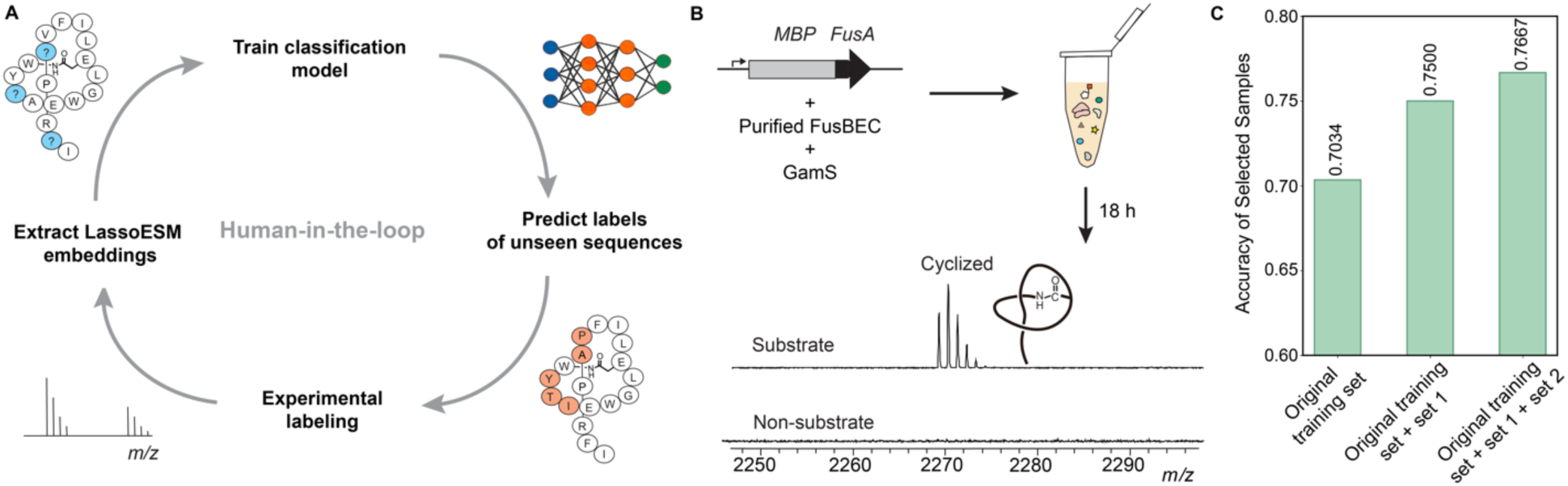
Experimental verification and optimization of model accuracy. **(A)** Human-in-the-loop workflow for optimizing the fusilassin cyclase substrate tolerance model. Embeddings for training data sequences were extracted from LassoESM. These embeddings were used to train a classifier, which then predicted the labels of unseen library sequences. Experimental validation was through CFB and MALDI-TOF-MS. The labeled data are then passed back into the workflow to improve model accuracy. **(B)** Experimental method to validate unseen sequences. The selected fusilassin variant sequences were produced using CFB and analyzed using MALDI-TOF-MS. **(C)** Model accuracy of the chosen samples across three rounds of experimental testing.

A library containing all 18 randomized sites on fusilassin has 20^18^ (2.6 × 10^23^) library members, which is too large to analyze reasonably. Therefore, two smaller libraries were selected for investigation. Library 1 varied positions 2, 3, 4, 13, and 14 (N = 3.2 million sequences), while library 2 varied positions 7, 8, 10, and 11 (N = 160,000 sequences). These positions were selected because they comprise a continuous surface on fusilassin, which could potentially engage with a target in future lasso peptide engineering campaigns (**Figure S3**). No sequences from either library were present in our initial training dataset.

The SVM model trained on LassoESM embeddings was used to classify all library sequences as substrates or non-substrates. A random selection of 118 fusilassin variant sequences (round 1) was experimentally validated (57 from library 1 and 61 from library 2) in CFB containing purified fusilassin biosynthetic enzymes. Lasso peptide cyclization was analyzed using MALDI-TOF-MS (**Figure 3B**). The overall model accuracy of round 1 was 70% (83/118) (**Tables S4-S7, Figures S4-S7**). This result underscored the extrapolation capability of the classification model with LassoESM embeddings, as the model performed well despite not having seen sequences in libraries 1 and 2.

To improve the model’s performance, we introduced a Human-in-the-Loop approach. The results from round 1 were incorporated into the initial training dataset. Embeddings for the new data were created with LassoESM, and the SVM classification model was retrained on all available data (**Figure 3A**). To verify the accuracy of the retrained model, 24 sequences were randomly selected from each library (round 2, N = 48) and evaluated using CFB. As anticipated, inclusion of round 1 data enhanced the model’s accuracy to 75% (36/48, **Figures 3C, S8-S9, Tables S8-S9**). In contrast, restricting the training data to the original dataset gave an accuracy of 69% (33/48).

The experimentally labeled data from rounds 1 and 2 were then incorporated into the training dataset, and the retrained SVM model was used to predict a more complex library (library 3), which varies positions 10 to 15 to encompass the entire fusilassin loop (library size of 64 million). A total of 30 fusilassin variant sequences were randomly selected from library 3 and evaluated using CFB. For these sequences, the model achieved an accuracy of 77% (23/30) (**Figures 3C, S10, Table S10**). The high accuracy further indicated that with the LassoESM embeddings, the trained SVM model demonstrated robust predictive accuracy even with sequences poorly represented in the initial training dataset. Table S3 summarizes the total number of sequences tested in CFB and the model accuracy for each library across all rounds.

### A general model for predicting lasso cyclase-lasso peptide pairs

LassoESM embeddings can also effectively predict more general lasso cyclase specificity tasks. For instance, diverse lasso peptides could be screened *in silico* or rationally engineered to bind a target of interest. However, extensive engineering may obscure which lasso cyclase can produce a desired lasso peptide. Thus, we designed a general model to predict any lasso cyclase-lasso peptide pair. A set of 6,599 unique lasso cyclase-lasso peptide pairs predicted via RODEO were used as training data.^10,13^ To create the negative samples, an equal number of non-natural cyclase-lasso peptide pairs were created by randomly mismatching cyclase and peptide pairs. Because most characterized lasso cyclases are specific for only one acceptor residue,^4^ the selected lasso peptides in the negative training set were verified to have a different acceptor residue from the native substrate (see Methods). The general model uses VanillaESM to extract embeddings for the lasso cyclase sequences and LassoESM to extract embeddings for the lasso peptide sequences. Rather than treating these two embeddings separately and concatenating them, a cross-attention layer was used to effectively capture interactions between lasso cyclases and lasso peptides (**Figure 4A**). The cross-attention layer emphasizes the amino acids in the lasso cyclase that likely interact with the lasso peptide substrate, thereby reducing noise and enhancing the model’s focus.

**Figure 4.**
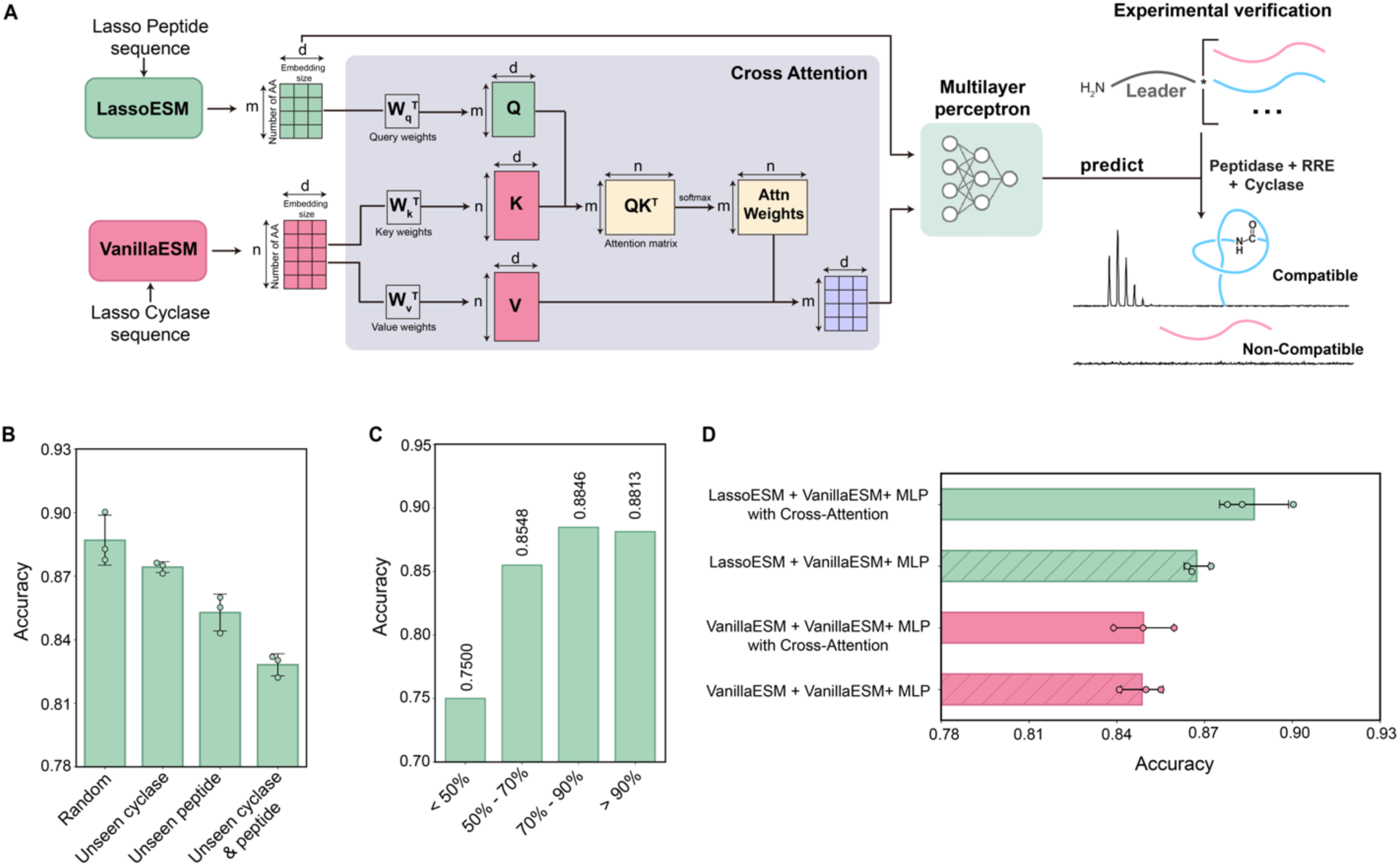
Architecture and evaluation of cyclase-peptide pair prediction. **(A)** The model architecture utilizes VanillaESM for cyclase embeddings and LassoESM for lasso peptide embeddings, incorporating a cross-attention layer to capture interactions between them. The attention-reweighted embeddings of lasso cyclases and peptides are averaged and then concatenated before being supplied to the MLP model to predict cyclase compatibility. The trained model is applied to predict the compatibility of other lasso peptides with FusC. **(B)** The model accuracy was evaluated on four dataset splits (random, unseen cyclase, unseen peptide, and unseen cyclase & peptide). (**C**) The model accuracy was assessed across different cyclase sequence similarities. The test set was divided into subsets based on maximum sequence similarity to those in the training set. Each bar represents the accuracy for cyclases within specific similarity ranges. **(D)** Ablation study evaluating the role of the cross-attention layer and LassoESM embeddings by accuracy scores of the general model. Error bars represent standard deviation of three repeats of model training.

The general model performance was evaluated under four different conditions: random, unseen cyclase, unseen peptide, and unseen cyclase & unseen peptide (**Figure 4B**). In the random condition, the dataset was randomly split into 70% for training, 15% for validation, and 15% for testing, resulting in an accuracy of 0.8870. The other three conditions simulate out-of-distribution scenarios, where the training and test sets have no overlapping cyclase or peptide sequences. In the most challenging condition with both unseen cyclase and unseen peptide, the model achieved an accuracy of 0.8281, demonstrating reliable and strong predictive performance for out-of-distribution data.

Naturally occurring lasso cyclases predicted by RODEO exhibit varying degrees of sequence similarity, with an average pairwise sequence identity of 42% across the dataset. To determine if model accuracy correlates with cyclase sequence similarity, we calculated the maximum pairwise sequence identity for each test cyclase relative to all cyclases in the training set. The results showed that model accuracy improves as the sequence identity of test cyclases increases compared to those in the training set (**Figure 4C**). For test cyclases with >50% sequence identity to any training cyclase, the accuracy exceeded 0.85. When sequence identity surpassed 70%, the model’s performance remained relatively stable.

Ablation analysis was conducted to evaluate the role of the important components in the model architecture. As shown in **Figure 4D**, removing the cross-attention layer decreased accuracy from 0.8870 to 0.8673, indicating the benefits of the cross-attention layer in enhancing model performance. In contrast to the model employing average pooling across all amino acids, the cross-attention layer enhances the model’s ability to focus on the amino acids that determine enzyme substrate specificity, thereby reducing the bias introduced by non-essential amino acids in the cyclase. Using VanillaESM to represent lasso peptides instead of LassoESM also resulted in a notable reduction in model accuracy. This demonstrates the effectiveness of LassoESM in predicting cyclase specificity for lasso peptides. We also observed that when the same language model was used to generate embeddings for both lasso cyclase and lasso peptides, the cross-attention layer failed to learn useful information, resulting in model performance that was no better than simple concatenation of the embeddings.

To evaluate LassoESM performance on experimentally verified cyclase-lasso peptide pairs, we used the trained model to predict the compatibility of FusC with other predicted naturally occurring lasso peptides. These sequences were tested using a chimeric engineering strategy, where the fusilassin leader peptide was fused to the selected core peptides. This enables the proper function of the RRE and leader peptidase, which are required for full cyclase activity. The set includes some chimeric peptides previously examined,^5^ which were selected based on cyclase similarity to FusC. It also contains Glu8- and Glu9-mer core peptides selected to maximize sequence diversity while maintaining the loop hydrophobicity previously shown to be preferred by FusC.^13^ Among the 35 experimentally validated chimeric peptide sequences, the model achieved an accuracy of 69%. Specifically, LassoESM correctly identified all three FusC-compatible lasso peptides and accurately classified 21 out of 32 non-compatible peptides (**Figure S11, Table S11**). Notably, the model predicted the FusC-compatible peptides with probabilities exceeding 0.7, indicating high confidence. Due to competition between the cyclase and endogenous proteases in the CFB, some of the 11 incorrectly predicted peptides may be poor substrates that would be detected in a pure setting.^5^ However, testing sequences by CFB is fast and more accurately reflects the heterologous expression conditions required for large scale lasso peptide production.

### Prediction of RNA polymerase inhibition by ubonodin and klebsidin variants

In addition to applying LassoESM embeddings to cyclase specificity prediction, we evaluated performance in an RNAP inhibition prediction task. Previous high-throughput screening studies explored the RNAP inhibitory activity of ubonodin variants and developed a deep learning model, DeepLasso, to predict RNAP inhibitory activity.^16^ VanillaESM, PeptideESM, and LassoESM were employed to extract embeddings for 8,885 literature-reported ubonodin variant sequences, and an MLP regressor was trained to predict the enrichment values, which act as estimates for RNAP inhibitory activity. The LassoESM embeddings model achieved the best performance (**Figure 5B-D**), with Pearson and Spearman correlation coefficients of 0.83 and 0.78, respectively. LassoESM obtained a mean absolute error (MAE) of 1.31, lower than that reported for DeepLasso (MAE of 2.20). DeepLasso uses lasso peptide sequence and topology information as input to train a model consisting of three convolutional neural network layers, two bidirectional long short-term memory layers, and one attention layer^16^. These results show that LassoESM embeddings yield a scenario where a simple two-layer MLP architecture resulted in superior model performance.

**Figure 5.**
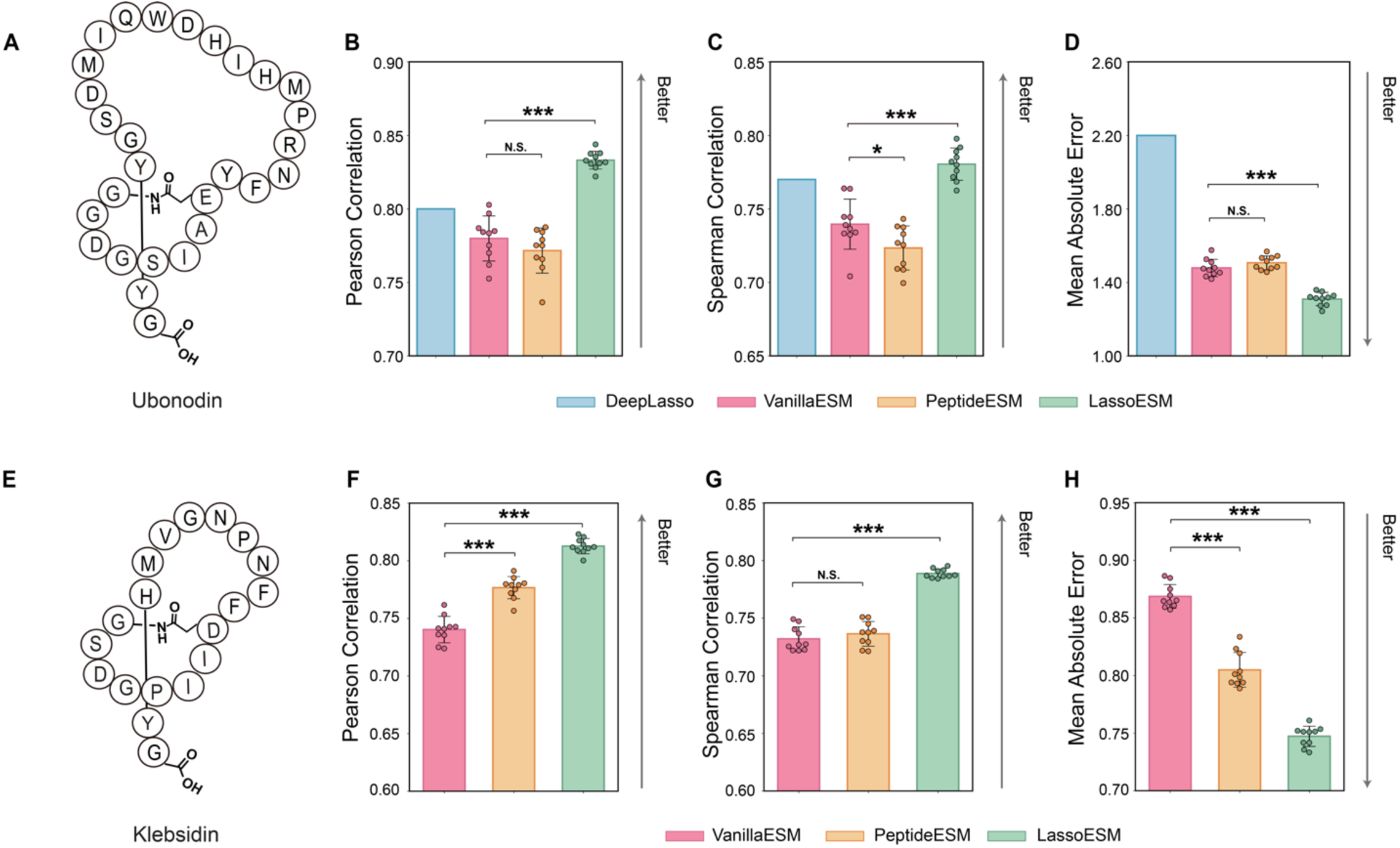
Comparison of RNA polymerase inhibitory activity predictions using different embeddings. **(A)** Bubble diagram of ubonodin. **(B-D)** Model performance of the MLP regressor for prediction of RNAP inhibition activity for the ubonodin variant dataset using embeddings from VanillaESM (pink), PeptideESM (orange), and LassoESM (green). Blue bars indicate the performance of DeepLasso.^16^ **(E)** Bubble diagram of klebsidin. **(F-H)** Model performance of the MLP regressor’s prediction of RNAP inhibition activity for the klebsidin variant dataset, using embeddings from VanillaESM (red), PeptideESM (orange), and LassoESM (green). Error bars: standard deviation calculated over 10 random seeds. *p*-value was calculated using a two-sided t-test. \**p* < 0.05, ** *p* < 0.01, *** *p* < 0.001. N.S., non-significant.

Similarly, we utilized LassoESM embeddings for another lasso peptide, klebsidin, to predict the RNAP inhibitory activity of its variants. We collected 340 klebsidin single-site variant sequences with enrichment values from a previous report.^42^ By searching for the best regression model, we trained an AdaBoost regressor on the klebsidin dataset (**Figure S12**).^42^ Even with the limited dataset, the LassoESM embeddings lead to Pearson and Spearman correlation coefficients of ∼0.80 and a MAE of ∼0.74. These results show significant improvement over those from VanillaESM embeddings, namely a 9% increase in Pearson correlation, 7% increase in Spearman correlation, and 14% decrease in MAE (**Figure 5F-H**).

### Conclusions

Lasso peptides are touted as promising scaffolds for drug development due to their remarkable stability and diverse biological activities. Many lasso peptide biosynthetic enzymes exhibit impressive substrate tolerance, allowing for the de novo design or rational engineering of novel lasso peptides with desired properties. However, predicting substrate tolerance and exploring sequence-activity relationships remains challenging due to the scarcity of experimentally labeled data. In this work, we utilized language models to improve the prediction of lasso peptide-related properties. To better capture lasso peptide features, we developed a lasso peptide-specific language model, LassoESM, which outperformed a generic PLM in a variety of downstream tasks, including predicting lasso cyclase substrate tolerance, identifying non-natural but compatible cyclase-substrate pairs, and predicting RNAP inhibition activity. Our results also show that embeddings from LassoESM enable accurate predictions even with small training sets, highlighting the potential of language models to address data scarcity and improve prediction accuracy. The pre-trained LassoESM offers optimal representations for lasso peptides and will be particularly useful for various lasso peptide-related tasks as high-throughput methods are developed for dataset generation.

Previous research has demonstrated that ESM-2 can recognize and store sequence motifs, which are specific patterns of amino acids that often appear together in proteins^53^. Since LassoESM was pre-trained on lasso peptide sequences, it is plausible that LassoESM learned to recognize the lasso fold pattern. Conversely, VanillaESM did not learn these specific sequence patterns because it was not supplied with the necessary sequences during training. Therefore, the underlying reason LassoESM outperforms VanillaESM in lasso peptide-related predictive tasks could be the tailored language model’s ability to learn the lasso fold.

While our computational workflow effectively characterized the substrate preference of lasso biosynthetic enzymes, the underlying molecular mechanisms that distinguish substrate from non-substrates remain unclear. Molecular dynamics simulations could offer valuable insights into these mechanisms by identifying key amino acids that influence substrate selectivity from both structural and dynamic perspectives. However, the lack of accurate structural knowledge of lasso biosynthetic enzymes and lasso peptides presents significant challenges in utilizing molecular dynamics simulations to understand these substrate preferences.

In summary, we developed a lasso peptide-tailored language model that provides effective representations for lasso peptides and enhances lasso peptide-related predictive tasks. Our efficient pipeline for characterizing the substrate preferences of lasso peptide biosynthetic enzymes and identifying lasso cyclase-lasso peptide pairs is a powerful tool for chemists aiming to rationally design functional and biocompatible lasso peptides for myriad biomedical and industrial applications.

## Acknowledgement

This work was supported by National Institutes of Health grants (R35GM142745 and R21AI167693 to D.S. and R01GM123998 to D.A.M.). S.E.B was supported by a National Science Foundation Graduate Research Fellowship (DGE 21-46756) and the University of Illinois Urbana-Champaign Illinois Distinguished Fellowship. The authors thank Joseph D. Clark for discussing LassoESM pretraining and Tanner J. Dean for discussing the cross-attention layer implementation. The authors also thank Soumajit Dutta and Song Yin for their valuable input during the preparation of this manuscript.

## Author information

### Authors and Affiliations

Center for Biophysics and Quantitative Biology, University of Illinois Urbana-Champaign, Urbana, IL, 61801

Xuenan Mi & Diwakar Shukla

Department of Chemistry, University of Illinois Urbana-Champaign, Urbana, IL, 61801

Susanna E. Barrett & Diwakar Shukla

Carl R. Woese Institute for Genomic Biology, University of Illinois Urbana-Champaign, Urbana, IL, 61801

Susanna E. Barrett

Department of Biochemistry, Vanderbilt University School of Medicine, and Department of Chemistry, Vanderbilt University, Nashville, TN 37232

Douglas A. Mitchell

Department of Bioengineering, Chemical and Biomolecular Engineering, University of Illinois Urbana-Champaign, Urbana, IL, 61801

Diwakar Shukla

Department of Bioengineering, University of Illinois Urbana-Champaign, Urbana, IL, 61801

Diwakar Shukla

## Supporting Information Available

Supplementary Tables 1-12 and Supplementary Figures 1–12

## Data Availability Statement

The code can be found at the following GitHub repository: https://github.com/ShuklaGroup/LassoESM

The LassoESM model can be downloaded from: https://huggingface.co/ShuklaGroupIllinois/LassoESM

## Ethics declarations

### Competing interests

S.E.B. and D.A.M. are inventors on a provisional patent application filed by the University of Illinois at Urbana-Champaign covering lasso cyclase engineering (US Provisional Application Ser. No. 63/673,853). D.A.M. is also a co-founder and owns stock in Lassogen, Inc. The other authors declare no competing interests.

## Notes

https://github.com/ShuklaGroup/LassoESM

https://huggingface.co/ShuklaGroupIllinois/LassoESM

