## Supplementary Information for "LassoESM: A tailored language model for enhanced lasso peptide property prediction"

**Table S1**. **Optimized hyperparameters of each classification model type trained on each set of embeddings for the fusilassin dataset.** RF: Random Forest; AdaBoost: Adaptive Boosting; SVM: Support Vector Machine; MLP: Multi-Layer Perceptron

| **Embedding** | **Downstream Model** | **Hyperparameters** |
|---|---|---|
| One-hot | RF | n_estimators = 100, max_depth = 20, max_features = "log2" |
| One-hot | AdaBoost | learning_rate = 1, n_estimators = 100 |
| One-hot | SVM | C = 1, kernel = "rbf" |
| One-hot | MLP | hidden_layer_sizes = 64, batch_size = 64, learning_rate = 0.01 |
| VanillaESM[1] | RF | n_estimators = 100, max_depth = 50, max_features = "sqrt" |
| VanillaESM | AdaBoost | learning_rate = 0.1, n_estimators = 200 |
| VanillaESM | SVM | C = 10, kernel = "linear" |
| VanillaESM | MLP | hidden_layer_sizes = (512, 64), batch_size = 16, learning_rate = 0.01 |
| PeptideESM[2] | RF | n_estimators = 100, max_depth = 10, max_features = "sqrt" |
| PeptideESM | AdaBoost | learning_rate = 0.1, n_estimators = 200 |
| PeptideESM | SVM | C = 10, kernel = "linear" |
| PeptideESM | MLP | hidden_layer_sizes = (256, 32), batch_size = 16, learning_rate = 0.001 |
| LassoESM | RF | n_estimators = 200, max_depth = 20, max_features = "sqrt" |
| LassoESM | AdaBoost | learning_rate = 0.1, n_estimators = 200 |
| LassoESM | SVM | C = 10, kernel = "linear" |
| LassoESM | MLP | hidden_layer_sizes = 256, batch_size = 16, learning_rate = 0.001 |

**Table S2. The optimal hyperparameters of each classification model type trained on each set of embeddings for the microcin J25 dataset.** RF: Random Forest; AdaBoost: Adaptive Boosting; SVM: Support Vector Machine; MLP: Multi-Layer Perceptron

| **Embedding** | **Downstream Model** | **Hyperparameters** |
|---|---|---|
| One-hot | RF | n_estimators = 100, max_depth = 50, max_features = "sqrt" |
| One-hot | AdaBoost | learning_rate = 5, n_estimators = 20 |
| One-hot | SVM | C = 0.1, kernel = "linear" |
| One-hot | MLP | hidden_layer_sizes = 64, batch_size = 16, learning_rate = 0.01 |
| VanillaESM[1] | RF | n_estimators = 20, max_depth = 10, max_features = "sqrt" |
| VanillaESM | AdaBoost | learning_rate = 10, n_estimators = 50 |
| VanillaESM | SVM | C = 10, kernel = "linear" |
| VanillaESM | MLP | hidden_layer_sizes = 256, batch_size = 16, learning_rate = 0.01 |
| PeptideESM[2] | RF | n_estimators = 20, max_depth = 50, max_features = "log2" |
| PeptideESM | AdaBoost | learning_rate = 1, n_estimators = 20 |
| PeptideESM | SVM | C = 10, kernel = "linear" |
| PeptideESM | MLP | hidden_layer_sizes = (256, 32), batch_size = 32, learning_rate = 0.001 |
| LassoESM | RF | n_estimators = 20, max_depth = 20, max_features = "log2" |
| LassoESM | AdaBoost | learning_rate = 1, n_estimators = 20 |
| LassoESM | SVM | C = 10, kernel = "linear" |
| LassoESM | MLP | hidden_layer_sizes = 256, batch_size = 32, learning_rate = 0.01 |

**Table S3. Summary of sequences verified in CFB to test the accuracy of the model.** The table includes the number of sequences tested for each library in each round of testing (set). It also contains the number of sequences with an accurate prediction and the percentage correct.

| Set | Library | Sequences tested in CFB | Number of sequences with accurate predictions |
|---|---|---|---|
| Round 1 | Library 1 | 57 | 38 (64%) |
| | Library 2 | 61 | 45 (73%) |
| Round 2 | Library 1 | 24 | 18 (75%) |
| | Library 2 | 24 | 18 (75%) |
| Round 3 | Library 3 | 30 | 23 (77%) |

**Table S4. Summary of round 1 fusilassin variants selected from library one and tested in CFB.** The sequence, model predicted substrate probability, model predicted label, and cyclized mass are listed. For the Predicted Label column, 0 indicates a predicted non-substrate, while 1 indicates a predicted substrate. The Substrate via CFB (cell-free biosynthesis) column indicates whether the sequence was a substrate, as determined experimentally by CFB and mass spectrometry (see Methods). N indicates the expected mass was not detected, while Y indicates the expected mass (i.e., the cyclized lasso peptide $[M+H]^+$) was detected. Red indicates that the model did not accurately predict the outcome (19 correct; 11 incorrect). Additional round 1 library one sequences are in **Table S5**. Spectra are shown in **Figure S4.** WT: Wild Type.

| **Library 1:** WXXXEWGLELIFXXPRFI | | | | |
|---|---|---|---|---|
| **FusA Variant Sequence** | **Predicted Probability** | **Predicted Label** | **Cyclized Mass $[M+H]^+$** | **Substrate via CFB?** |
| WYTAEWGLELIFVFPRFI | WT | WT | WT | Y |
| WNEEEWGLELIFTEPRFI | 0.0065 | 0 | 2290.1 | N |
| WKAVEWGLELIFKLPRFI | 0.0085 | 0 | 2227.3 | N |
| WIENEWGLELIFVDPRFI | 0.0303 | 0 | 2258.2 | N |
| WMAKEWGLELIFHDPRFI | 0.0514 | 0 | 2270.2 | N |
| WALIEWGLELIFNWPRFI | 0.0799 | 0 | 2285.2 | N |
| WNLVEWGLELIFDIPRFI | 0.0818 | 0 | 2242.2 | N |
| WIESEWGLELIFVEPRFI | 0.0870 | 0 | 2245.2 | N |
| WMWHEWGLELIFWLPRFI | 0.0886 | 0 | 2441.2 | N |
| WMADEWGLELIFTNPRFI | 0.1175 | 0 | 2220.1 | N |
| WQIVEWGLELIFGTPRFI | 0.1192 | 0 | 2186.2 | N |
| WTNTEWGLELIFRFPRFI | 0.1499 | 0 | 2307.2 | Y |
| WFPNEWGLELIFHVPRFI | 0.1963 | 0 | 2282.2 | Y |
| WIETEWGLELIFVQPRFI | 0.1995 | 0 | 2258.2 | N |
| WKIAEWGLELIFALPRFI | 0.2202 | 0 | 2184.2 | Y |
| WRANEWGLELIFGMPRFI | 0.2235 | 0 | 2217.1 | Y |
| WNYFEWGLELIFYCPRFI | 0.3086 | 0 | 2378.1 | Y |
| WQLAEWGLELIFNIPRFI | 0.3116 | 0 | 2227.2 | N |
| WMLKEWGLELIFTAPRFI | 0.3222 | 0 | 2232.2 | N |
| WLVKEWGLELIFAFPRFI | 0.3632 | 0 | 2246.2 | N |
| WMNQEWGLELIFVQPRFI | 0.3717 | 0 | 2288.2 | N |
| WMMEEWGLELIFSMPRFI | 0.4341 | 0 | 2297.1 | Y |
| WFHMEWGLELIFNAPRFI | 0.4605 | 0 | 2281.1 | Y |
| WFTEEWGLELIFMYPRFI | 0.4847 | 0 | 2359.2 | Y |
| WLMMEWGLELIFHLPRFI | 0.4946 | 0 | 2313.2 | Y |
| WTQIEWGLELIFAFPRFI | 0.5718 | 1 | 2248.2 | Y |
| WYSAEWGLELIFVGPRFI | 0.5905 | 1 | 2165.1 | Y |
| WWANEWGLELIFFPPRFI | 0.6012 | 1 | 2303.2 | N |
| WYPAEWGLELIFTFPRFI | 0.6084 | 1 | 2267.2 | Y |
| WYTIEWGLELIFPAPRFI | 0.7343 | 1 | 2233.2 | Y |
| WSFTEWGLELIFVPPRFI | 0.7414 | 1 | 2219.2 | N |

**Table S5. Summary of round 1 fusilassin variants for library one tested using CFB cont.** The sequence, model predicted substrate probability, model predicted label, and cyclized mass are listed. For the Predicted Label column, 0 indicates a predicted non-substrate, while 1 indicates a predicted substrate. The Substrate via CFB (cell-free biosynthesis) column indicates whether the sequence was a substrate, as determined experimentally by CFB and mass spectrometry (see Methods). N indicates the expected mass was not detected while Y indicates the expected mass (i.e., the cyclized lasso peptide $[M+H]^+$) was detected. Red indicates that the model did not accurately predict the outcome (19 correct; 8 incorrect). Additional round 1 library one sequences are in **Table S4**. Spectra are shown in **Figure S5.** WT: Wild Type.

| **Library 1:** WXXXEWGLELIFXXPRFI | | | | |
|---|---|---|---|---|
| **Sequence** | **Predicted Probability** | **Model Prediction** | **Cyclized Mass $[M+H]^+$** | **Substrate in CFB ?** |
| WYTAEWGLELIFVFPRFI | WT | WT | WT | Y |
| WPCEEWGLELIFTKPRFI | 0.0095 | 0 | 2246.1 | N |
| WCSPEWGLELIFEKPRFI | 0.0201 | 0 | 2232.1 | N |
| WDEAEWGLELIFCCPRFI | 0.0338 | 0 | 2209.0 | Y |
| WFPVEWGLELIFWSPRFI | 0.0461 | 0 | 2304.2 | N |
| WPDTEWGLELIFYFPRFI | 0.0447 | 0 | 2311.2 | Y |
| WRADEWGLELIFQMPRFI | 0.0601 | 0 | 2289.2 | Y |
| WEEAEWGLELIFCFPRFI | 0.0769 | 0 | 2267.1 | Y |
| WGFLEWGLELIFNWPRFI | 0.0879 | 0 | 2305.2 | N |
| WPEYEWGLELIFCTPRFI | 0.0987 | 0 | 2281.1 | N |
| WTADEWGLELIFQYPRFI | 0.0998 | 0 | 2266.1 | N |
| WPYAEWGLELIFTQPRFI | 0.1170 | 0 | 2248.2 | N |
| WASDEWGLELIFTVPRFI | 0.1322 | 0 | 2161.1 | N |
| WGCLEWGLELIFADPRFI | 0.1412 | 0 | 2147.1 | Y |
| WFVREWGLELIFTHPRFI | 0.1486 | 0 | 2328.2 | N |
| WWWGEWGLELIFGIPRFI | 0.1848 | 0 | 2287.2 | N |
| WCNVEWGLELIFPRPRFI | 0.1899 | 0 | 2257.2 | N |
| WCPDEWGLELIFTFPRFI | 0.1964 | 0 | 2251.1 | Y |
| WPSFEWGLELIFVHPRFI | 0.2135 | 0 | 2255.2 | N |
| WDHAEWGLELIFAIPRFI | 0.2355 | 0 | 2195.1 | N |
| WQSPEWGLELIFFVPRFI | 0.2534 | 0 | 2246.2 | N |
| WECAEWGLELIFCAPRFI | 0.2587 | 0 | 2165.0 | Y |
| WHHTEWGLELIFGPPRFI | 0.3541 | 0 | 2217.1 | N |
| WASCEWGLELIFDYPRFI | 0.3853 | 0 | 2227.1 | Y |
| WMMMEWGLELIFMMPRFI | 0.5000 | 1 | 2343.1 | Y |
| WFHMEWGLELIFGAPRFI | 0.5642 | 1 | 2231.1 | Y |
| WQNMEWGLELIFALPRFI | 0.6466 | 1 | 2245.2 | Y |
| WYATEWGLELIFNMPRFI | 0.7325 | 1 | 2268.1 | Y |

**Table S6. Summary of round 1 fusilassin variants selected from library two and tested in CFB.** The sequence, model predicted substrate probability, model predicted label, and cyclized mass are listed. For the Predicted Label column, 0 indicates a predicted non-substrate, while 1 indicates a predicted substrate. The Substrate via CFB (cell-free biosynthesis) column indicates whether the sequence was a substrate, as determined experimentally by CFB and mass spectrometry (see Methods). N indicates the expected mass was not detected while Y indicates the expected mass (i.e., the cyclized lasso peptide $[M+H]^+$) was detected. Red indicates that the model did not accurately predict the outcome (18 correct; 9 incorrect). Additional round 1 library two sequences are in **Table S7**. Spectra are shown in **Figure S6.** WT: Wild Type.

| **Library 2:** WYTAEWXXEXXFVFPRFI | | | | |
|---|---|---|---|---|
| **Sequence** | **Predicted Probability** | **Predicted Label** | **Cyclized Mass $[M+H]^+$** | **Substrate in CFB?** |
| WYTAEWGLELIFVFPRFI | WT | WT | WT | Y |
| WYTAEWNGEKYFVFPRFI | 0.0085 | 0 | 2335.1 | N |
| WYTAEWWEENAFVFPRFI | 0.0937 | 0 | 2373.1 | N |
| WYTAEWCDESQFVFPRFI | 0.1806 | 0 | 2306.0 | N |
| WYTAEWAAEDAFVFPRFI | 0.2084 | 0 | 2201.0 | N |
| WYTAEWDPETAFVFPRFI | 0.2386 | 0 | 2257.1 | N |
| WYTAEWYMEQGFVFPRFI | 0.2687 | 0 | 2352.1 | N |
| WYTAEWAGEKTFVFPRFI | 0.2747 | 0 | 2230.1 | N |
| WYTAEWGGERVFVFPRFI | 0.2853 | 0 | 2242.1 | N |
| WYTAEWDDETMFVFPRFI | 0.2897 | 0 | 2335.0 | N |
| WYTAEWKFETTFVFPRFI | 0.3438 | 0 | 2350.2 | N |
| WYTAEWAQEDYFVFPRFI | 0.3537 | 0 | 2350.1 | N |
| WYTAEWCCEKAFVFPRFI | 0.4009 | 0 | 2278.1 | N |
| WYTAEWEGETYFVFPRFI | 0.4245 | 0 | 2323.1 | N |
| WYTAEWCHEAQFVFPRFI | 0.4618 | 0 | 2312.1 | N |
| WYTAEWASEDYFVFPRFI | 0.4709 | 0 | 2309.1 | N |
| WYTAEWEQEFFFVFPRFI | 0.6334 | 1 | 2424.1 | N |
| WYTAEWVVEIHFVFPRFI | 0.6606 | 1 | 2321.2 | N |
| WYTAEWPEELFFVFPRFI | 0.6755 | 1 | 2359.2 | N |
| WYTAEWPHELYFVFPRFI | 0.6918 | 1 | 2383.2 | N |
| WYTAEWPFECIFVFPRFI | 0.7093 | 1 | 2333.1 | N |
| WYTAEWPLEMGFVFPRFI | 0.7170 | 1 | 2271.1 | N |
| WYTAEWGLEETFVFPRFI | 0.7780 | 1 | 2273.1 | N |
| WYTAEWVPEAIFVFPRFI | 0.8568 | 1 | 2253.1 | N |
| WYTAEWIPEVYFVFPRFI | 0.8806 | 1 | 2345.2 | N |
| WYTAEWGEELLFVFPRFI | 0.9084 | 1 | 2285.1 | Y |
| WYTAEWCTESVFVFPRFI | 0.9416 | 1 | 2263.1 | Y |
| WYTAEWATELMFVFPRFI | 0.9500 | 1 | 2289.1 | Y |

**Table S7. Summary of round 1 fusilassin variants for library two tested using CFB cont.** The sequence, model predicted substrate probability, model predicted label, and cyclized mass are listed. For the Predicted Label column, 0 indicates a predicted non-substrate, while 1 indicates a predicted substrate. The Substrate via CFB (cell-free biosynthesis) column indicates whether the sequence was a substrate, as determined experimentally by CFB and mass spectrometry (see Methods). N indicates the expected mass was not detected, while Y indicates the expected mass (i.e., the cyclized lasso peptide $[M+H]^+$) was detected. Red indicates that the model did not accurately predict the outcome (27 correct; 7 incorrect). Additional round 1 library two sequences are in **Table S6**. Spectra are shown in **Figure S7.** WT: Wild Type.

| **Library 2:** WYTAEWXXEXXFVFPRFI | | | | |
|---|---|---|---|---|
| **Sequence** | **Predicted Probability** | **Model Prediction** | **Cyclized Mass $[M+H]^+$** | **Substrate in CFB ?** |
| WYTAEWGLELIFVFPRFI | WT | WT | WT | Y |
| WYTAEWEREKKFVFPRFI | 0.0025 | 0 | 2414.2 | N |
| WYTAEWIREHRFVFPRFI | 0.0037 | 0 | 2435.2 | N |
| WYTAEWEWEKRFVFPRFI | 0.0111 | 0 | 2472.2 | N |
| WYTAEWSSEKKFVFPRFI | 0.0156 | 0 | 2303.2 | N |
| WYTAEWNWEHRFVFPRFI | 0.0284 | 0 | 2466.2 | N |
| WYTAEWDMEIKFVFPRFI | 0.0366 | 0 | 2360.1 | N |
| WYTAEWPYERHFVFPRFI | 0.0423 | 0 | 2426.2 | N |
| WYTAEWINEGGFVFPRFI | 0.0545 | 0 | 2214.1 | N |
| WYTAEWIMENIFVFPRFI | 0.1208 | 0 | 2344.2 | N |
| WYTAEWQLEGGFVFPRFI | 0.1962 | 0 | 2228.1 | N |
| WYTAEWGLESKFVFPRFI | 0.2138 | 0 | 2258.1 | N |
| WYTAEWGLENEFVFPRFI | 0.3233 | 0 | 2286.1 | Y |
| WYTAEWWFEEFFVFPRFI | 0.3635 | 0 | 2482.2 | N |
| WYTAEWSLEGGFVFPRFI | 0.3974 | 0 | 2187.1 | N |
| WYTAEWELEGIFVFPRFI | 0.4026 | 0 | 2285.1 | N |
| WYTAEWGKEPFFVFPRFI | 0.4281 | 0 | 2302.1 | Y |
| WYTAEWILEGSFVFPRFI | 0.4524 | 0 | 2243.1 | N |
| WYTAEWKLELGFVFPRFI | 0.4604 | 0 | 2284.2 | N |
| WYTAEWFFELEFVFPRFI | 0.4899 | 0 | 2409.2 | N |
| WYTAEWGSEMQFVFPRFI | 0.5276 | 1 | 2276.1 | Y |
| WYTAEWEWELLFVFPRFI | 0.5563 | 1 | 2414.2 | N |
| WYTAEWRNEMIFVFPRFI | 0.6202 | 1 | 2387.2 | Y |
| WYTAEWGPEAAFVFPRFI | 0.6206 | 1 | 2169.1 | Y |
| WYTAEWMMELMFVFPRFI | 0.6308 | 1 | 2379.1 | Y |
| WYTAEWKVETIFVFPRFI | 0.7209 | 1 | 2314.2 | Y |
| WYTAEWGGESMFVFPRFI | 0.7223 | 1 | 2205.0 | Y |
| WYTAEWGIEDLFVFPRFI | 0.7506 | 1 | 2271.1 | Y |
| WYTAEWTCEFCFVFPRFI | 0.8035 | 1 | 2327.0 | N |
| WYTAEWSAEWVFVFPRFI | 0.8046 | 1 | 2316.1 | N |
| WYTAEWMTEGFFVFPRFI | 0.8165 | 1 | 2309.1 | Y |
| WYTAEWGDELVFVFPRFI | 0.8345 | 1 | 2257.1 | Y |

| | | | | |
|---|---|---|---|---|
| WYTAEW**HS**E**LC**FVFPRFI | 0.8430 | 1 | 2313.1 | N |
| WYTAEW**YS**E**CL**FVFPRFI | 0.8540 | 1 | 2339.1 | Y |
| WYTAEW**MF**E**SF**FVFPRFI | 0.9203 | 1 | 2385.1 | N |

**Table S8. Summary of randomly selected round 2 fusilassin variants for library one tested using CFB.** The sequence, model predicted substrate probability, model predicted label, and cyclized mass are listed. For the Predicted Label column, 0 indicates a predicted non-substrate, while 1 indicates a predicted substrate. The Substrate via CFB (cell-free biosynthesis) column indicates whether the sequence was a substrate, as determined experimentally by CFB and mass spectrometry (see Methods). N indicates the expected mass was not detected while Y indicates the expected mass (i.e., the cyclized lasso peptide $[M+H]^+$) was detected. Red indicates that the model did not accurately predict the outcome (18 correct; 6 incorrect). Spectra are shown in **Figure S8.** WT: Wild Type.

| **Library 1:** WXXXEWGLELIFXXPRFI | | | | |
|---|---|---|---|---|
| **Sequence** | **Predicted Probability** | **Model Prediction** | **Cyclized Mass $[M+H]^+$** | **Substrate in CFB ?** |
| WYTAEWGLELIFVFPRFI | WT | WT | WT | Y |
| WQREEWGLELIFKRPRFI | 0.0012 | 0 | 2384.3 | N |
| WPETEWGLELIFWWPRFI | 0.0106 | 0 | 2386.2 | N |
| WAHEEWGLELIFWHPRFI | 0.0295 | 0 | 2347.2 | N |
| WEVCEWGLELIFRLPRFI | 0.0406 | 0 | 2287.2 | N |
| WNHKEWGLELIFSEPRFI | 0.0821 | 0 | 2282.2 | Y |
| WFNHEWGLELIFRSPRFI | 0.0973 | 0 | 2328.2 | N |
| WRPGEWGLELIFYHPRFI | 0.1017 | 0 | 2297.2 | N |
| WYFYEWGLELIFMKPRFI | 0.1874 | 0 | 2419.2 | N |
| WMFCEWGLELIFFWPRFI | 0.2466 | 0 | 2401.2 | Y |
| WCDGEWGLELIFGVPRFI | 0.2596 | 0 | 2118.0 | N |
| WMIQEWGLELIFEFPRFI | 0.3106 | 0 | 2335.2 | Y |
| WFIDEWGLELIFATPRFI | 0.3599 | 0 | 2234.2 | Y |
| WNWREWGLELIFVMPRFI | 0.3821 | 0 | 2373.2 | N |
| WCMIEWGLELIFVWPRFI | 0.3933 | 0 | 2319.2 | N |
| WLASEWGLELIFEIPRFI | 0.4331 | 0 | 2200.2 | N |
| WYHGEWGLELIFHIPRFI | 0.5068 | 1 | 2294.2 | Y |
| WAWTEWGLELIFMTPRFI | 0.6358 | 1 | 2277.1 | Y |
| WYTNEWGLELIFNLPRFI | 0.6433 | 1 | 2292.2 | Y |
| WLFMEWGLELIFMLPRFI | 0.6549 | 1 | 2322.2 | Y |
| WLYGEWGLELIFNYPRFI | 0.7105 | 1 | 2297.2 | Y |
| WHALEWGLELIFCMPRFI | 0.7184 | 1 | 2242.1 | Y |
| WMANEWGLELIFSCPRFI | 0.8451 | 1 | 2193.1 | N |
| WYTYEWGLELIFLTPRFI | 0.8458 | 1 | 2328.2 | Y |
| WFCMEWGLELIFGPPRFI | 0.8820 | 1 | 2222.1 | N |

**Table S9. Summary of randomly selected round 2 fusilassin variants for library two tested using CFB.** The sequence, model predicted substrate probability, model predicted label, and cyclized mass are listed. For the Predicted Label column, 0 indicates a predicted non-substrate, while 1 indicates a predicted substrate. The Substrate via CFB (cell-free biosynthesis) column indicates whether the sequence was a substrate, as determined experimentally by CFB and mass spectrometry (see Methods). N indicates the expected mass was not detected, while Y indicates the expected mass (i.e., the cyclized lasso peptide $[M+H]^+$) was detected. Red indicates that the model did not accurately predict the outcome (18 correct; 6 incorrect). Spectra are shown in **Figure S9.** WT: Wild Type.

| **Library 2:** WYTAEWXXEXXFVFPRFI | | | | |
|---|---|---|---|---|
| **Sequence** | **Predicted Probability** | **Model Prediction** | **Cyclized Mass $[M+H]^+$** | **Substrate in CFB ?** |
| WYTAEWGLELIFVFPRFI | WT | WT | WT | Y |
| WYTAEWEFERKFVFPRFI | 0.0029 | 0 | 2432.2 | N |
| WYTAEWKPEADFVFPRFI | 0.0261 | 0 | 2283.1 | N |
| WYTAEWICEWNFVFPRFI | 0.0297 | 0 | 2388.1 | N |
| WYTAEWHTEPQFVFPRFI | 0.0319 | 0 | 2335.1 | N |
| WYTAEWCSERQFVFPRFI | 0.0523 | 0 | 2346.1 | N |
| WYTAEWNVESNFVFPRFI | 0.1224 | 0 | 2286.1 | Y |
| WYTAEWMYEKLFVFPRFI | 0.1461 | 0 | 2407.2 | N |
| WYTAEWFGEWAFVFPRFI | 0.2178 | 0 | 2333.1 | N |
| WYTAEWIIELFFVFPRFI | 0.2213 | 0 | 2358.2 | N |
| WYTAEWAPESTFVFPRFI | 0.2241 | 0 | 2228.1 | N |
| WYTAEWSQEMQFVFPRFI | 0.2277 | 0 | 2346.1 | N |
| WYTAEWWQEMVFVFPRFI | 0.3888 | 0 | 2416.2 | Y |
| WYTAEWGTEINFVFPRFI | 0.4008 | 0 | 2257.1 | Y |
| WYTAEWLMEQLFVFPRFI | 0.4140 | 0 | 2357.2 | Y |
| WYTAEWGAEEAFVFPRFI | 0.4657 | 0 | 2200.0 | Y |
| WYTAEWAVESAFVFPRFI | 0.6358 | 1 | 2200.1 | Y |
| WYTAEWDMEAVFVFPRFI | 0.6726 | 1 | 2288.1 | Y |
| WYTAEWCLESIFVFPRFI | 0.7293 | 1 | 2288.1 | Y |
| WYTAEWGIELGFVFPRFI | 0.7389 | 1 | 2212.1 | Y |
| WYTAEWAEEVIFVFPRFI | 0.7717 | 1 | 2284.1 | N |
| WYTAEWGNEVIFVFPRFI | 0.8603 | 1 | 2255.1 | Y |
| WYTAEWGWEVMFVFPRFI | 0.8708 | 1 | 2345.1 | Y |
| WYTAEWALELVFVFPRFI | 0.8731 | 1 | 2268.2 | Y |
| WYTAEWAAEVFFVFPRFI | 0.9174 | 1 | 2260.1 | Y |

**Table S10. Summary of randomly selected round 3 fusilassin variants for library three tested using CFB.** The sequence, model predicted substrate probability, model predicted label, and cyclized mass are listed. For the Predicted Label column, 0 indicates a predicted non-substrate, while 1 indicates a predicted substrate. The Substrate via CFB (cell-free biosynthesis) column indicates whether the sequence was a substrate, as determined experimentally by CFB and mass spectrometry (see Methods). N indicates the expected mass was not detected, while Y indicates the expected mass (i.e., the cyclized lasso peptide [M+H]$^+$) was detected. Red indicates that the model did not accurately predict the outcome (23 correct; 7 incorrect). Spectra are shown in **Figure S10.** WT: Wild Type.

| **Library 3:** WYTAEWGLEXXXXXXRFI | | | | |
|---|---|---|---|---|
| **Sequence** | **Predicted Probability** | **Model Prediction** | **Cyclized [M+H]$^+$** | **substrate in CFB?** |
| WYTAEWGLELIFVFPRFI | WT | WT | WT | Y |
| WYTAEWGLE**FKPKEE**RFI | 0.0008 | 0 | 2311.1 | N |
| WYTAEWGLE**NWRMEI**RFI | 0.0043 | 0 | 2382.1 | N |
| WYTAEWGLE**KQCNMM**RFI | 0.0115 | 0 | 2288.0 | N |
| WYTAEWGLE**TKCCAY**RFI | 0.0133 | 0 | 2222.0 | N |
| WYTAEWGLE**GTKWFT**RFI | 0.0206 | 0 | 2273.1 | N |
| WYTAEWGLE**ECIQAR**RFI | 0.0319 | 0 | 2253.1 | N |
| WYTAEWGLE**QTAMDY**RFI | 0.0579 | 0 | 2262.0 | N |
| WYTAEWGLE**GWRNTT**RFI | 0.0586 | 0 | 2268.1 | N |
| WYTAEWGLE**SLHPYI**RFI | 0.0940 | 0 | 2263.1 | N |
| WYTAEWGLE**TYGVGG**RFI | 0.1174 | 0 | 2087.0 | N |
| WYTAEWGLE**RWMMMG**RFI | 0.1224 | 0 | 2345.1 | N |
| WYTAEWGLE**TLEFPR**RFI | 0.1230 | 0 | 2296.1 | N |
| WYTAEWGLE**WGNTFN**RFI | 0.1249 | 0 | 2272.1 | N |
| WYTAEWGLE**LYGTCD**RFI | 0.1642 | 0 | 2205.0 | N |
| WYTAEWGLE**TNPPCF**RFI | 0.3410 | 0 | 2212.0 | N |
| WYTAEWGLE**MWDVLP**RFI | 0.5282 | 1 | 2294.1 | Y |
| WYTAEWGLE**AFVLHA**RFI | 0.5490 | 1 | 2191.1 | N |
| WYTAEWGLE**PLFMPT**RFI | 0.5540 | 1 | 2239.1 | N |
| WYTAEWGLE**IFWGIN**RFI | 0.5765 | 1 | 2283.1 | N |
| WYTAEWGLE**LITLLN**RFI | 0.5963 | 1 | 2220.2 | Y |
| WYTAEWGLE**YLFFCQ**RFI | 0.6018 | 1 | 2354.1 | N |
| WYTAEWGLE**MYWNVP**RFI | 0.6110 | 1 | 2343.1 | Y |
| WYTAEWGLE**TAMAFG**RFI | 0.7124 | 1 | 2131.0 | N |
| WYTAEWGLE**LWFCSP**RFI | 0.7325 | 1 | 2286.1 | Y |
| WYTAEWGLE**VFGLLT**RFI | 0.7347 | 1 | 2183.1 | Y |
| WYTAEWGLE**YLFYMT**RFI | 0.7618 | 1 | 2371.1 | N |
| WYTAEWGLE**MGVAFC**RFI | 0.7641 | 1 | 2161.0 | N |
| WYTAEWGLE**MFLFMS**RFI | 0.8049 | 1 | 2309.1 | Y |
| WYTAEWGLE**LNFFCP**RFI | 0.8545 | 1 | 2274.1 | Y |
| WYTAEWGLE**TLMYAP**RFI | 0.9157 | 1 | 2229.1 | Y |

**Table S11. Summary of chimeric sequences tested using CFB.** The chimeric sequences contained the FusA leader peptide and a naturally occurring core peptide identified from the RODEO mined dataset of all lasso peptides. Core peptides were selected based on sequence diversity, hydrophobicity, and acceptor identity and position. The sequence name, sequence, model predicted substrate probability, model predicted label, and cyclized mass $[M+H]^+$ are listed. For the model prediction label column, a 0 means the sequence is predicted to be a non-substrate, while a 1 means it is predicted to be a substrate. The last column indicates if the sequence was a substrate determined experimentally via CFB where N means no peak was detected with MALDI-TOF-MS and Y means a peak with the mass of the cyclized lasso $[M+H]^+$ was detected. Orange indicates that the model accurately predicted the outcome. [3]Sequences previously published are indicated with the reference in the "substrate in CFB?" column and were selected for testing based on the high sequence identity of their cognate cyclase to FusC. Spectra are shown in **Figure S12.**

| Name | Sequence | Predicted Probability | Model Prediction | Cyclized $[M+H]^+$ | substrate in CFB? |
|---|---|---|---|---|---|
| SegA | GMPGAFVEILGEDDKPAGLTQE | 0.0000 | 0 | 2256.1 | N |
| Sma6A | GSIGALGDENGLHKQVGISND | 0.0000 | 0 | 2063.0 | N |
| ChloA | ASMNEIAPELVGDKTQRFGG | 0.0005 | 0 | 2102.0 | N |
| KorA | SGSLTPTESMTMMMRKP | 0.0015 | 0 | 1866.9 | N |
| BbaA | DALPGPYLEMGIFPSRTVEG | 0.0027 | 0 | 2131.0 | N |
| MflA | GDIYPGVESLDPNSRIN | 0.0036 | 0 | 1827.9 | N |
| OliA | GEMGTLLEFILERDS | 0.0041 | 0 | 1691.8 | N |
| KunA | GNPDGDEVEIDEQLYFKP | 0.0042 | 0 | 2046.9 | N |
| BacA | AGSKGYQEVVMILMDWKV | 0.0140 | 0 | 2036.0 | N |
| SidA | GYFVGSYKEWIVRRIV | 0.0154 | 0 | 1954.1 | N |
| SulA | SGALLDLVEVFLVSHNS | 0.0200 | 0 | 1781.9 | N |
| FlaA | GGDGPGTEMLTFHYDPG | 0.0283 | 0 | 1732.7 | N |
| BulA | ADSPDGEFELIPVQRWTMTGVRFH | 0.0314 | 0 | 2770.3 | N |
| CacA | GYPLGAQEIVGFLSRDQ | 0.0472 | 0 | 1831.9 | N |
| PpcA | APAMFGSPEIGNLIYAYRV | 0.0935 | 0 | 2051.0 | N |
| RhoA | GGGIWWVEWVGKFN | 0.1172 | 0 | 1616.8 | N |
| Cau31A | SFDVGTIKEGLVSQYYFA | 0.1569 | 0 | 2006.0 | N |
| MobA | YIGLEGSEPITHSFSKFW | 0.2144 | 0 | 2080.0 | N[3] |
| CarA | GYFYGSYKEFLSRRIV | 0.2294 | 0 | 1967.0 | N |
| SruA | RGGEPIWEEVVVPWDYWV | 0.2686 | 0 | 2198.1 | N[3] |
| EnsA | GNNSAGVHETLGPKRYRTF | 0.3175 | 0 | 2086.0 | N |
| JesA | TGPKTFTQEILTLFTPS | 0.5959 | 1 | 1863.0 | N |
| NetA | LGISGGPELLLLKRM | 0.6598 | 1 | 1578.9 | N |
| EndA | WQQGRGFEVLFFLPRMI | 0.7096 | 1 | 2106.1 | Y |
| HalA | YKSGRGLELWLFLPRMV | 0.8026 | 1 | 2047.1 | Y[3] |
| NbsA | YFGLTGYENVIHFYDRL | 0.8632 | 1 | 2089.0 | N[3] |
| CelA | WIQGKWGLEIYLIFPRYL | 0.8708 | 1 | 2277.3 | Y[3] |
| MthA | YNAINKLEIIFIWPRLFN | 0.8751 | 1 | 2246.2 | N[3] |

| | | | | | |
|---|---|---|---|---|---|
| XylA | VFYVRNGEEVLWFFDTWI | 0.8879 | 1 | 2302.1 | N |
| HaiA | GLPWTRTEALYGYKIT | 0.8919 | 1 | 1851.0 | N |
| SleA | LYGVRNDEEINWHFDYWT | 0.8987 | 1 | 2339.0 | N[3] |
| NcaA | YVGLRNRESLLGYPRNIW | 0.9058 | 1 | 2188.2 | N[3] |
| RubA | ALGLHGAEPFFPTLHTSWW | 0.9061 | 1 | 2149.1 | N[3] |
| RegA | LGTHRGPERVLPARFSL | 0.9085 | 1 | 1888.1 | N |
| SalA | FIGPIHYEGILLWHSWF | 0.9248 | 1 | 2097.1 | N |

**Table S12. Primers and ultramer sequences used in this study.** From left to right, 5' to 3'. F means forward primer while R means reverse primer. Names for library primers indicate the variant sequence at the variable positions in the library.

| | |
|---|---|
| CFB_F | GCTATCATGCCATACCGCGAAAGGTTTTGCGCCATTCG |
| CFB_R | AACCGTCTATCAGGGCGATGGCCCACTACGTGAACCATC |
| **Library 1 (WXXXEWGLELIFXXPRFI)** | |
| RAN_GM_F | GGGCCTCGAGCTGATCTTCGGAATGCCGCGCTTCATCTGAGCG |
| RAN_GM_R | GAAGATCAGCTCGAGGCCCCATTCATTAGCACGCCAGCCGGTGGCCTC |
| YAT_NM_F | GGGCCTCGAGCTGATCTTCAATATGCCGCGCTTCATCTGAGCG |
| YAT_NM_R | GAAGATCAGCTCGAGGCCCCATTCAGTTGCGTACCAGCCGGTGGCCTC |
| MMM_MM_F | GGGCCTCGAGCTGATCTTCATGATGCCGCGCTTCATCTGAGCG |
| MMM_MM_R | GAAGATCAGCTCGAGGCCCCATTCCATCATCATCCAGCCGGTGGCCTC |
| FHM_GA_F | GGGCCTCGAGCTGATCTTCGGAGCCCCGCGCTTCATCTGAGCG |
| FHM_GA_R | GAAGATCAGCTCGAGGCCCCATTCCATGTGAAACCAGCCGGTGGCCTC |
| IET_VQ_F | GGGCCTCGAGCTGATCTTCGTTCAACCGCGCTTCATCTGAGCG |
| IET_VQ_R | GAAGATCAGCTCGAGGCCCCATTCAGTTTCGATCCAGCCGGTGGCCTC |
| IEN_VD_F | GGGCCTCGAGCTGATCTTCGTTGACCCGCGCTTCATCTGAGCG |
| IEN_VD_R | GAAGATCAGCTCGAGGCCCCATTCATTTTCAATCCAGCCGGTGGCCTC |
| FHM_NA_F | GGGCCTCGAGCTGATCTTCAACGCACCGCGCTTCATCTGAGCG |
| FHM_NA_R | GAAGATCAGCTCGAGGCCCCATTCCATATGGAACCAGCCGGTGGCCTC |
| CNV_PR_F | GGGCCTCGAGCTGATCTTCCCACGCCCGCGCTTCATCTGAGCG |
| CNV_PR_R | GAAGATCAGCTCGAGGCCCCATTCCACGTTGCACCAGCCGGTGGCCTC |
| PSF_VH_F | GGGCCTCGAGCTGATCTTCGTGCATCCGCGCTTCATCTGAGCG |
| PSF_VH_R | GAAGATCAGCTCGAGGCCCCATTCAAACGACGGCCAGCCGGTGGCCTC |
| FVR_TH_F | GGGCCTCGAGCTGATCTTCACTCACCCGCGCTTCATCTGAGCG |
| FVR_TH_R | GAAGATCAGCTCGAGGCCCCATTCGCGTACAAACCAGCCGGTGGCCTC |
| PCE_TK_F | GGGCCTCGAGCTGATCTTCACAAAGCCGCGCTTCATCTGAGCG |
| PCE_TK_R | GAAGATCAGCTCGAGGCCCCATTCCTCGCAAGGCCAGCCGGTGGCCTC |
| CSP_EK_F | GGGCCTCGAGCTGATCTTCGAGAAGCCGCGCTTCATCTGAGCG |
| CSP_EK_R | GAAGATCAGCTCGAGGCCCCATTCGGGCGAACACCAGCCGGTGGCCTC |
| FPV_WS_F | GGGCCTCGAGCTGATCTTCTGGAGTCCGCGCTTCATCTGAGCG |
| FPV_WS_R | GAAGATCAGCTCGAGGCCCCATTCGACAGGAAACCAGCCGGTGGCCTC |
| ASC_DY_F | GGGCCTCGAGCTGATCTTCGATTATCCGCGCTTCATCTGAGCG |
| ASC_DY_R | GAAGATCAGCTCGAGGCCCCATTCACAGCTCGCCCAGCCGGTGGCCTC |
| PEY_CT_F | GGGCCTCGAGCTGATCTTCTGCACACCGCGCTTCATCTGAGCG |
| PEY_CT_R | GAAGATCAGCTCGAGGCCCCATTCGTACTCCGGCCAGCCGGTGGCCTC |
| CPD_TF_F | GGGCCTCGAGCTGATCTTCACCTTTCCGCGCTTCATCTGAGCG |
| CPD_TF_R | GAAGATCAGCTCGAGGCCCCATTCGTCCGGGCACCAGCCGGTGGCCTC |
| ALI_NW_F | GGGCCTCGAGCTGATCTTCAATTGGCCGCGCTTCATCTGAGCG |

| | |
|---|---|
| ALI_NW_R | GAAGATCAGCTCGAGGCCCCATTCAATCAGTGCCCAGCCGGTGGCCTC |
| FPN_HV_F | GGGCCTCGAGCTGATCTTCCACGTACCGCGCTTCATCTGAGCG |
| FPN_HV_R | GAAGATCAGCTCGAGGCCCCATTCGTTGGGGAACCAGCCGGTGGCCTC |
| IES_VE_F | GGGCCTCGAGCTGATCTTCGTGGAGCCGCGCTTCATCTGAGCG |
| IES_VE_R | GAAGATCAGCTCGAGGCCCCATTCACTCTCGATCCAGCCGGTGGCCTC |
| KIA_AL_F | GGGCCTCGAGCTGATCTTCGCATTGCCGCGCTTCATCTGAGCG |
| KIA_AL_R | GAAGATCAGCTCGAGGCCCCATTCGGCGATTTTCCAGCCGGTGGCCTC |
| KAV_KL_F | GGGCCTCGAGCTGATCTTCAAGTTGCCGCGCTTCATCTGAGCG |
| KAV_KL_R | GAAGATCAGCTCGAGGCCCCATTCAACCGCCTTCCAGCCGGTGGCCTC |
| MAK_HD_F | GGGCCTCGAGCTGATCTTCCATGACCCGCGCTTCATCTGAGCG |
| MAK_HD_R | GAAGATCAGCTCGAGGCCCCATTCCTTGGCCATCCAGCCGGTGGCCTC |
| MAD_TN_F | GGGCCTCGAGCTGATCTTCACTAATCCGCGCTTCATCTGAGCG |
| MAD_TN_R | GAAGATCAGCTCGAGGCCCCATTCGTCCGCCATCCAGCCGGTGGCCTC |
| NLV_DI_F | GGGCCTCGAGCTGATCTTCGACATTCCGCGCTTCATCTGAGCG |
| NLV_DI_R | GAAGATCAGCTCGAGGCCCCATTCGACTAAGTTCCAGCCGGTGGCCTC |
| QIV_GT_F | GGGCCTCGAGCTGATCTTCGGTACGCCGCGCTTCATCTGAGCG |
| QIV_GT_R | GAAGATCAGCTCGAGGCCCCATTCTACAATTTGCCAGCCGGTGGCCTC |
| QLA_NI_F | GGGCCTCGAGCTGATCTTCAACATCCCGCGCTTCATCTGAGCG |
| QLA_NI_R | GAAGATCAGCTCGAGGCCCCATTCCGCCAGTTGCCAGCCGGTGGCCTC |
| YTI_PA_F | GGGCCTCGAGCTGATCTTCCCCGCGCCGCGCTTCATCTGAGCG |
| YTI_PA_R | GAAGATCAGCTCGAGGCCCCATTCAATTGTATACCAGCCGGTGGCCTC |
| YPA_TF_F | GGGCCTCGAGCTGATCTTCACATTTCCGCGCTTCATCTGAGCG |
| YPA_TF_R | GAAGATCAGCTCGAGGCCCCATTCGGCAGGATACCAGCCGGTGGCCTC |
| TQI_AF_F | GGGCCTCGAGCTGATCTTCGCCTTTCCGCGCTTCATCTGAGCG |
| TQI_AF_R | GAAGATCAGCTCGAGGCCCCATTCAATCTGGGTCCAGCCGGTGGCCTC |
| PDT_YF_F | GGGCCTCGAGCTGATCTTCTATTTTCCGCGCTTCATCTGAGCG |
| PDT_YF_R | GAAGATCAGCTCGAGGCCCCATTCCGTGTCCGGCCAGCCGGTGGCCTC |
| YSA_VG_F | GGGCCTCGAGCTGATCTTCGTTGGCCCGCGCTTCATCTGAGCG |
| YSA_VG_R | GAAGATCAGCTCGAGGCCCCATTCGGCTGAGTACCAGCCGGTGGCCTC |
| SFT_VP_F | GGGCCTCGAGCTGATCTTCGTGCCGCCGCGCTTCATCTGAGCG |
| SFT_VP_R | GAAGATCAGCTCGAGGCCCCATTCTGTGAAGGACCAGCCGGTGGCCTC |
| PYA_TQ_F | GGGCCTCGAGCTGATCTTCACACAGCCGCGCTTCATCTGAGCG |
| PYA_TQ_R | GAAGATCAGCTCGAGGCCCCATTCAGCGTAGGGCCAGCCGGTGGCCTC |
| ASD_TV_F | GGGCCTCGAGCTGATCTTCACTGTACCGCGCTTCATCTGAGCG |
| ASD_TV_R | GAAGATCAGCTCGAGGCCCCATTCGTCTGAGGCCCAGCCGGTGGCCTC |
| QSP_FV_F | GGGCCTCGAGCTGATCTTCTTCGTGCCGCGCTTCATCTGAGCG |
| QSP_FV_R | GAAGATCAGCTCGAGGCCCCATTCCGGGGATTGCCAGCCGGTGGCCTC |
| TAD_QY_F | GGGCCTCGAGCTGATCTTCCAATATCCGCGCTTCATCTGAGCG |
| TAD_QY_R | GAAGATCAGCTCGAGGCCCCATTCGTCTGCGGTCCAGCCGGTGGCCTC |
| MLK_TA_F | GGGCCTCGAGCTGATCTTCACTGCCCCGCGCTTCATCTGAGCG |
| MLK_TA_R | GAAGATCAGCTCGAGGCCCCATTCTTTTAACATCCAGCCGGTGGCCTC |

| | |
|---|---|
| NYF_YC_F | GGGCCTCGAGCTGATCTTCTACTGCCCGCGCTTCATCTGAGCG |
| NYF_YC_R | GAAGATCAGCTCGAGGCCCCATTCAAAATAGTTCCAGCCGGTGGCCTC |
| NEE_TE_F | GGGCCTCGAGCTGATCTTCACTGAGCCGCGCTTCATCTGAGCG |
| NEE_TE_R | GAAGATCAGCTCGAGGCCCCATTCTTCCTCATTCCAGCCGGTGGCCTC |
| LVK_AF_F | GGGCCTCGAGCTGATCTTCGCGTTCCCGCGCTTCATCTGAGCG |
| LVK_AF_R | GAAGATCAGCTCGAGGCCCCATTCTTTTACTAACCAGCCGGTGGCCTC |
| TNT_RF_F | GGGCCTCGAGCTGATCTTCCGCTTCCCGCGCTTCATCTGAGCG |
| TNT_RF_R | GAAGATCAGCTCGAGGCCCCATTCGGTATTGGTCCAGCCGGTGGCCTC |
| LMM_HL_F | GGGCCTCGAGCTGATCTTCCACCTTCCGCGCTTCATCTGAGCG |
| LMM_HL_R | GAAGATCAGCTCGAGGCCCCATTCCATCATTAACCAGCCGGTGGCCTC |
| MWH_WL_F | GGGCCTCGAGCTGATCTTCTGGCTTCCGCGCTTCATCTGAGCG |
| MWH_WL_R | GAAGATCAGCTCGAGGCCCCATTCATGCCACATCCAGCCGGTGGCCTC |
| QNM_AL_F | GGGCCTCGAGCTGATCTTCGCACTTCCGCGCTTCATCTGAGCG |
| QNM_AL_R | GAAGATCAGCTCGAGGCCCCATTCCATATTTTGCCAGCCGGTGGCCTC |
| MME_SM_F | GGGCCTCGAGCTGATCTTCTCTATGCCGCGCTTCATCTGAGCG |
| MME_SM_R | GAAGATCAGCTCGAGGCCCCATTCCTCCATCATCCAGCCGGTGGCCTC |
| MNQ_VQ_F | GGGCCTCGAGCTGATCTTCGTTCAGCCGCGCTTCATCTGAGCG |
| MNQ_VQ_R | GAAGATCAGCTCGAGGCCCCATTCCTGGTTCATCCAGCCGGTGGCCTC |
| FTE_MY_F | GGGCCTCGAGCTGATCTTCATGTACCCGCGCTTCATCTGAGCG |
| FTE_MY_R | GAAGATCAGCTCGAGGCCCCATTCCTCAGTGAACCAGCCGGTGGCCTC |
| ECA_CA_F | GGGCCTCGAGCTGATCTTCTGTGCTCCGCGCTTCATCTGAGCG |
| ECA_CA_R | GAAGATCAGCTCGAGGCCCCATTCAGCGCATTCCCAGCCGGTGGCCTC |
| DEA_CC_F | GGGCCTCGAGCTGATCTTCTGCTGTCCGCGCTTCATCTGAGCG |
| DEA_CC_R | GAAGATCAGCTCGAGGCCCCATTCAGCCTCGTCCCAGCCGGTGGCCTC |
| GCL_AD_F | GGGCCTCGAGCTGATCTTCGCAGATCCGCGCTTCATCTGAGCG |
| GCL_AD_R | GAAGATCAGCTCGAGGCCCCATTCAAGGCACCCCCAGCCGGTGGCCTC |
| EEA_CF_F | GGGCCTCGAGCTGATCTTCTGCTTCCCGCGCTTCATCTGAGCG |
| EEA_CF_R | GAAGATCAGCTCGAGGCCCCATTCAGCCTCTTCCCAGCCGGTGGCCTC |
| YPA_WF_F | GGGCCTCGAGCTGATCTTCTGGTTTCCGCGCTTCATCTGAGCG |
| YPA_WF_R | GAAGATCAGCTCGAGGCCCCATTCGGCTGGGTACCAGCCGGTGGCCTC |
| DHA_AI_F | GGGCCTCGAGCTGATCTTCGCGATTCCGCGCTTCATCTGAGCG |
| DHA_AI_R | GAAGATCAGCTCGAGGCCCCATTCTGCGTGATCCCAGCCGGTGGCCTC |
| WWG_GI_F | GGGCCTCGAGCTGATCTTCGGGATCCCGCGCTTCATCTGAGCG |
| WWG_GI_R | GAAGATCAGCTCGAGGCCCCATTCACCCCACCACCAGCCGGTGGCCTC |
| RAD_QM_F | GGGCCTCGAGCTGATCTTCCAAATGCCGCGCTTCATCTGAGCG |
| RAD_QM_R | GAAGATCAGCTCGAGGCCCCATTCATCCGCGCGCCAGCCGGTGGCCTC |
| HHT_GP_F | GGGCCTCGAGCTGATCTTCGGACCGCCGCGCTTCATCTGAGCG |
| HHT_GP_R | GAAGATCAGCTCGAGGCCCCATTCGGTATGATGCCAGCCGGTGGCCTC |
| WAN_FP_F | GGGCCTCGAGCTGATCTTCTTCCCTCCGCGCTTCATCTGAGCG |
| WAN_FP_R | GAAGATCAGCTCGAGGCCCCATTCGTTTGCCCACCAGCCGGTGGCCTC |
| GFL_NW_F | GGGCCTCGAGCTGATCTTCAATTGGCCGCGCTTCATCTGAGCG |

| | |
|---|---|
| GFL_NW_R | GAAGATCAGCTCGAGGCCCCATTCTAAGAAACCCCAGCCGGTGGCCTC |
| PET_WW_F | GGGCCTCGAGCTGATCTTCTGGTGGCCGCGCTTCATCTGAGCG |
| PET_WW_R | GAAGATCAGCTCGAGGCCCCATTCGGTTTCTGGCCAGCCGGTGGCCTC |
| QRE_KR_F | GGGCCTCGAGCTGATCTTCAAGCGTCCGCGCTTCATCTGAGCG |
| QRE_KR_R | GAAGATCAGCTCGAGGCCCCATTCTTCGCGTTGCCAGCCGGTGGCCTC |
| AHE_WH_F | GGGCCTCGAGCTGATCTTCTGGCATCCGCGCTTCATCTGAGCG |
| AHE_WH_R | GAAGATCAGCTCGAGGCCCCATTCTTCGTGGGCCCAGCCGGTGGCCTC |
| EVC_RL_F | GGGCCTCGAGCTGATCTTCCGCTTGCCGCGCTTCATCTGAGCG |
| EVC_RL_R | GAAGATCAGCTCGAGGCCCCATTCGCAGACTTCCCAGCCGGTGGCCTC |
| RPG_YH_F | GGGCCTCGAGCTGATCTTCTATCACCCGCGCTTCATCTGAGCG |
| RPG_YH_R | GAAGATCAGCTCGAGGCCCCATTCCCCTGGGCGCCAGCCGGTGGCCTC |
| NWR_VM_F | GGGCCTCGAGCTGATCTTCGTAATGCCGCGCTTCATCTGAGCG |
| NWR_VM_R | GAAGATCAGCTCGAGGCCCCATTCGCGCCAATTCCAGCCGGTGGCCTC |
| CMI_VW_F | GGGCCTCGAGCTGATCTTCGTCTGGCCGCGCTTCATCTGAGCG |
| CMI_VW_R | GAAGATCAGCTCGAGGCCCCATTCAATCATACACCAGCCGGTGGCCTC |
| FNH_RS_F | GGGCCTCGAGCTGATCTTCCGCTCCCCGCGCTTCATCTGAGCG |
| FNH_RS_R | GAAGATCAGCTCGAGGCCCCATTCGTGGTTGAACCAGCCGGTGGCCTC |
| CDG_GV_F | GGGCCTCGAGCTGATCTTCGGTGTGCCGCGCTTCATCTGAGCG |
| CDG_GV_R | GAAGATCAGCTCGAGGCCCCATTCCCCGTCACACCAGCCGGTGGCCTC |
| NHK_SE_F | GGGCCTCGAGCTGATCTTCAGTGAACCGCGCTTCATCTGAGCG |
| NHK_SE_R | GAAGATCAGCTCGAGGCCCCATTCCTTATGGTTCCAGCCGGTGGCCTC |
| YFY_MK_F | GGGCCTCGAGCTGATCTTCATGAAACCGCGCTTCATCTGAGCG |
| YFY_MK_R | GAAGATCAGCTCGAGGCCCCATTCGTAAAAATACCAGCCGGTGGCCTC |
| FCM_GP_F | GGGCCTCGAGCTGATCTTCGGACCCCCGCGCTTCATCTGAGCG |
| FCM_GP_R | GAAGATCAGCTCGAGGCCCCATTCCATGCAAAACCAGCCGGTGGCCTC |
| AWT_MT_F | GGGCCTCGAGCTGATCTTCATGACTCCGCGCTTCATCTGAGCG |
| AWT_MT_R | GAAGATCAGCTCGAGGCCCCATTCGGTCCATGCCCAGCCGGTGGCCTC |
| MIQ_EF_F | GGGCCTCGAGCTGATCTTCGAGTTTCCGCGCTTCATCTGAGCG |
| MIQ_EF_R | GAAGATCAGCTCGAGGCCCCATTCCTGAATCATCCAGCCGGTGGCCTC |
| MFC_FW_F | GGGCCTCGAGCTGATCTTCTTTTGGCCGCGCTTCATCTGAGCG |
| MFC_FW_R | GAAGATCAGCTCGAGGCCCCATTCACAAAACATCCAGCCGGTGGCCTC |
| FID_AT_F | GGGCCTCGAGCTGATCTTCGCTACACCGCGCTTCATCTGAGCG |
| FID_AT_R | GAAGATCAGCTCGAGGCCCCATTCGTCGATAAACCAGCCGGTGGCCTC |
| HAL_CM_F | GGGCCTCGAGCTGATCTTCTGCATGCCGCGCTTCATCTGAGCG |
| HAL_CM_R | GAAGATCAGCTCGAGGCCCCATTCCAGTGCATGCCAGCCGGTGGCCTC |
| YHG_HI_F | GGGCCTCGAGCTGATCTTCCATATTCCGCGCTTCATCTGAGCG |
| YHG_HI_R | GAAGATCAGCTCGAGGCCCCATTCGCCATGGTACCAGCCGGTGGCCTC |
| LAS_EI_F | GGGCCTCGAGCTGATCTTCGAAATTCCGCGCTTCATCTGAGCG |
| LAS_EI_R | GAAGATCAGCTCGAGGCCCCATTCACTAGCTAACCAGCCGGTGGCCTC |
| YTY_LT_F | GGGCCTCGAGCTGATCTTCCTGACCCCGCGCTTCATCTGAGCG |
| YTY_LT_R | GAAGATCAGCTCGAGGCCCCATTCGTAAGTATACCAGCCGGTGGCCTC |

| | |
|---|---|
| LFM_ML_F | GGGCCTCGAGCTGATCTTCATGCTGCCGCGCTTCATCTGAGCG |
| LFM_ML_R | GAAGATCAGCTCGAGGCCCCATTCCATAAACAGCCAGCCGGTGGCCTC |
| MAN_SC_F | GGGCCTCGAGCTGATCTTCTCTTGCCCGCGCTTCATCTGAGCG |
| MAN_SC_R | GAAGATCAGCTCGAGGCCCCATTCGTTTGCCATCCAGCCGGTGGCCTC |
| LYG_NY_F | GGGCCTCGAGCTGATCTTCAACTATCCGCGCTTCATCTGAGCG |
| LYG_NY_R | GAAGATCAGCTCGAGGCCCCATTCTCCATAAAGCCAGCCGGTGGCCTC |
| YTN_NL_F | GGGCCTCGAGCTGATCTTCAATCTGCCGCGCTTCATCTGAGCG |
| YTN_NL_R | GAAGATCAGCTCGAGGCCCCATTCATTGGTGTACCAGCCGGTGGCCTC |
| **Library 2 (WYTAEWXXEXXFVFPRFI)** | |
| CTESV_F | TGTACGGAGAGTGTTTTCGTCTTCCCGCGCTTCA |
| CTESV_R | GACGAAAACACTCTCCGTACACCATTCCGCGGTGTACC |
| MMELM_F | ATGATGGAGCTGATGTTCGTCTTCCCGCGCTTCA |
| MMELM_R | GACGAACATCAGCTCCATCATCCATTCCGCGGTGTACC |
| MFESF_F | ATGTTTGAGAGTTTTTTCGTCTTCCCGCGCTTCA |
| MFESF_R | GACGAAAAAACTCTCAAACATCCATTCCGCGGTGTACC |
| DPETA_F | GACCCAGAGACGGCATTCGTCTTCCCGCGCTTCA |
| DPETA_R | GACGAATGCCGTCTCTGGGTCCCATTCCGCGGTGTACC |
| ATELM_F | GCCACTGAGTTAATGTTCGTCTTCCCGCGCTTCA |
| ATELM_R | GACGAACATTAACTCAGTGGCCCATTCCGCGGTGTACC |
| KVETI_F | AAAGTTGAGACAATCTTCGTCTTCCCGCGCTTCA |
| KVETI_R | GACGAAGATTGTCTCAACTTTCCATTCCGCGGTGTACC |
| GGESM_F | GGTGGTGAGTCTATGTTCGTCTTCCCGCGCTTCA |
| GGESM_R | GACGAACATAGACTCACCACCCCATTCCGCGGTGTACC |
| HSELC_F | CATTCGGAGTTATGTTTCGTCTTCCCGCGCTTCA |
| HSELC_R | GACGAAACATAACTCCGAATGCCATTCCGCGGTGTACC |
| KFETT_F | AAATTTGAGACCACGTTCGTCTTCCCGCGCTTCA |
| KFETT_R | GACGAACGTGGTCTCAAATTTCCATTCCGCGGTGTACC |
| EWELL_F | GAATGGGAGCTTCTTTTCGTCTTCCCGCGCTTCA |
| EWELL_R | GACGAAAAGAAGCTCCCATTCCCATTCCGCGGTGTACC |
| NWEHR_F | AATTGGGAGCATCGTTTCGTCTTCCCGCGCTTCA |
| NWEHR_R | GACGAAACGATGCTCCCAATTCCATTCCGCGGTGTACC |
| IREHR_F | ATCCGCGAGCACCGCTTCGTCTTCCCGCGCTTCA |
| IREHR_R | GACGAAGCGGTGCTCGCGGATCCATTCCGCGGTGTACC |
| PYERH_F | CCCTATGAGCGCCACTTCGTCTTCCCGCGCTTCA |
| PYERH_R | GACGAAGTGGCGCTCATAGGGCCATTCCGCGGTGTACC |
| EWEKR_F | GAATGGGAGAAACGCTTCGTCTTCCCGCGCTTCA |
| EWEKR_R | GACGAAGCGTTTCTCCCATTCCCATTCCGCGGTGTACC |
| EREKK_F | GAGCGTGAGAAAAAATTCGTCTTCCCGCGCTTCA |
| EREKK_R | GACGAATTTTTTCTCACGCTCCCATTCCGCGGTGTACC |
| SSEKK_F | TCCTCGGAGAAAAAGTTCGTCTTCCCGCGCTTCA |

| | |
|---|---|
| SSEKK_R | GACGAACTTTTTCTCCGAGGACCATTCCGCGGTGTACC |
| DMEIK_F | GACATGGAGATCAAGTTCGTCTTCCCGCGCTTCA |
| DMEIK_R | GACGAACTTGATCTCCATGTCCCATTCCGCGGTGTACC |
| VVEIH_F | GTGGTCGAGATTCATTTCGTCTTCCCGCGCTTCA |
| VVEIH_R | GACGAAATGAATCTCGACCACCCATTCCGCGGTGTACC |
| QLEGG_F | CAATTAGAGGGCGGGTTCGTCTTCCCGCGCTTCA |
| QLEGG_R | GACGAACCCGCCCTCTAATTGCCATTCCGCGGTGTACC |
| INEGG_F | ATCAACGAGGGAGGGTTCGTCTTCCCGCGCTTCA |
| INEGG_R | GACGAACCCTCCCTCGTTGATCCATTCCGCGGTGTACC |
| CHEAQ_F | TGCCATGAAGCCCAATTCGTCTTCCCGCGCTTCA |
| CHEAQ_R | GACGAATTGGGCTTCATGGCACCATTCCGCGGTGTACC |
| GPEAA_F | GGGCCGGAGGCGGCATTCGTCTTCCCGCGCTTCA |
| GPEAA_R | GACGAATGCCGCCTCCGGCCCCCATTCCGCGGTGTACC |
| AQEDY_F | GCCCAGGAAGACTACTTCGTCTTCCCGCGCTTCA |
| AQEDY_R | GACGAAGTAGTCTTCCTGGGCCCATTCCGCGGTGTACC |
| ASEDY_F | GCTTCAGAGGATTACTTCGTCTTCCCGCGCTTCA |
| ASEDY_R | GACGAAGTAATCCTCTGAAGCCCATTCCGCGGTGTACC |
| AGEKT_F | GCAGGAGAGAAAACGTTCGTCTTCCCGCGCTTCA |
| AGEKT_R | GACGAACGTTTTCTCTCCTGCCCATTCCGCGGTGTACC |
| GDELV_F | GGCGATGAGTTGGTCTTCGTCTTCCCGCGCTTCA |
| GDELV_R | GACGAAGACCAACTCATCGCCCCATTCCGCGGTGTACC |
| WEENA_F | TGGGAGGAAAATGCTTTCGTCTTCCCGCGCTTCA |
| WEENA_R | GACGAAAGCATTTTCCTCCCACCATTCCGCGGTGTACC |
| YMEQG_F | TATATGGAGCAGGGCTTCGTCTTCCCGCGCTTCA |
| YMEQG_R | GACGAAGCCCTGCTCCATATACCATTCCGCGGTGTACC |
| CDESQ_F | TGCGACGAGTCGCAGTTCGTCTTCCCGCGCTTCA |
| CDESQ_R | GACGAACTGCGACTCGTCGCACCATTCCGCGGTGTACC |
| DDETM_F | GATGACGAGACTATGTTCGTCTTCCCGCGCTTCA |
| DDETM_R | GACGAACATAGTCTCGTCATCCCATTCCGCGGTGTACC |
| EGETY_F | GAGGGAGAAACATATTTCGTCTTCCCGCGCTTCA |
| EGETY_R | GACGAAATATGTTTCTCCCTCCCATTCCGCGGTGTACC |
| GLEET_F | GGGCTTGAAGAGACTTTCGTCTTCCCGCGCTTCA |
| GLEET_R | GACGAAAGTCTCTTCAAGCCCCCATTCCGCGGTGTACC |
| KLELG_F | AAACTGGAATTGGGGTTCGTCTTCCCGCGCTTCA |
| KLELG_R | GACGAACCCCAATTCCAGTTTCCATTCCGCGGTGTACC |
| SLEGG_F | TCACTTGAAGGCGGATTCGTCTTCCCGCGCTTCA |
| SLEGG_R | GACGAATCCGCCTTCAAGTGACCATTCCGCGGTGTACC |
| ELEGI_F | GAGTTAGAGGGTATTTTCGTCTTCCCGCGCTTCA |
| ELEGI_R | GACGAAAATACCCTCTAACTCCCATTCCGCGGTGTACC |
| ILEGS_F | ATTTTAGAAGGTAGTTTCGTCTTCCCGCGCTTCA |
| ILEGS_R | GACGAAACTACCTTCTAAAATCCATTCCGCGGTGTACC |

| | |
|---|---|
| GEELL_F | GGTGAAGAGTTATTGTTCGTCTTCCCGCGCTTCA |
| GEELL_R | GACGAACAATAACTCTTCACCCCATTCCGCGGTGTACC |
| PLEMG_F | CCCCTGGAAATGGGGTTCGTCTTCCCGCGCTTCA |
| PLEMG_R | GACGAACCCCATTTCCAGGGGCCATTCCGCGGTGTACC |
| GLENE_F | GGGCTGGAAAACGAATTCGTCTTCCCGCGCTTCA |
| GLENE_R | GACGAATTCGTTTTCCAGCCCCCATTCCGCGGTGTACC |
| GLESK_F | GGCTTGGAGTCGAAGTTCGTCTTCCCGCGCTTCA |
| GLESK_R | GACGAACTTCGACTCCAAGCCCCATTCCGCGGTGTACC |
| AAEDA_F | GCAGCAGAAGATGCATTCGTCTTCCCGCGCTTCA |
| AAEDA_R | GACGAATGCATCTTCTGCTGCCCATTCCGCGGTGTACC |
| CCEKA_F | TGCTGTGAGAAGGCTTTCGTCTTCCCGCGCTTCA |
| CCEKA_R | GACGAAAGCCTTCTCACAGCACCATTCCGCGGTGTACC |
| GGERV_F | GGAGGCGAGCGCGTGTTCGTCTTCCCGCGCTTCA |
| GGERV_R | GACGAACACGCGCTCGCCTCCCCATTCCGCGGTGTACC |
| GIEDL_F | GGGATCGAGGACTTATTCGTCTTCCCGCGCTTCA |
| GIEDL_R | GACGAATAAGTCCTCGATCCCCCATTCCGCGGTGTACC |
| GSEMQ_F | GGTTCGGAAATGCAATTCGTCTTCCCGCGCTTCA |
| GSEMQ_R | GACGAATTGCATTTCCGAACCCCATTCCGCGGTGTACC |
| MTEGF_F | ATGACAGAGGGTTTTTTCGTCTTCCCGCGCTTCA |
| MTEGF_R | GACGAAAAAACCCTCTGTCATCCATTCCGCGGTGTACC |
| NGEKY_F | AACGGTGAGAAATACTTCGTCTTCCCGCGCTTCA |
| NGEKY_R | GACGAAGTATTTCTCACCGTTCCATTCCGCGGTGTACC |
| RNEMI_F | CGCAACGAGATGATTTTCGTCTTCCCGCGCTTCA |
| RNEMI_R | GACGAAAATCATCTCGTTGCGCCATTCCGCGGTGTACC |
| SAEWV_F | TCTGCTGAGTGGGTCTTCGTCTTCCCGCGCTTCA |
| SAEWV_R | GACGAAGACCCACTCAGCAGACCATTCCGCGGTGTACC |
| TCEFC_F | ACATGTGAGTTCTGCTTCGTCTTCCCGCGCTTCA |
| TCEFC_R | GACGAAGCAGAACTCACATGTCCATTCCGCGGTGTACC |
| YSECL_F | TATTCAGAATGCTTATTCGTCTTCCCGCGCTTCA |
| YSECL_R | GACGAATAAGCATTCTGAATACCATTCCGCGGTGTACC |
| EQEFF_F | GAACAGGAATTCTTCTTCGTCTTCCCGCGCTTCA |
| EQEFF_R | GACGAAGAAGAATTCCTGTTCCCATTCCGCGGTGTACC |
| FFELE_F | TTTTTCGAGTTAGAGTTCGTCTTCCCGCGCTTCA |
| FFELE_R | GACGAACTCTAACTCGAAAAACCATTCCGCGGTGTACC |
| GKEPF_F | GGAAAGGAGCCATTTTTCGTCTTCCCGCGCTTCA |
| GKEPF_R | GACGAAAAATGGCTCCTTTCCCCATTCCGCGGTGTACC |
| IMENI_F | ATCATGGAAAACATTTTCGTCTTCCCGCGCTTCA |
| IMENI_R | GACGAAAATGTTTTCCATGATCCATTCCGCGGTGTACC |
| IPEVY_F | ATTCCAGAGGTCTACTTCGTCTTCCCGCGCTTCA |
| IPEVY_R | GACGAAGTAGACCTCTGGAATCCATTCCGCGGTGTACC |
| PEELF_F | CCGGAGGAGCTTTTTTTCGTCTTCCCGCGCTTCA |

| | |
|---|---|
| PEELF_R | GACGAAAAAAAGCTCCTCCGGCCATTCCGCGGTGTACC |
| PFECI_F | CCGTTTGAATGTATTTTCGTCTTCCCGCGCTTCA |
| PFECI_R | GACGAAAATACATTCAAACGGCCATTCCGCGGTGTACC |
| PHELY_F | CCCCACGAGTTATATTTCGTCTTCCCGCGCTTCA |
| PHELY_R | GACGAAATATAACTCGTGGGGCCATTCCGCGGTGTACC |
| VPEAI_F | GTTCCCGAAGCTATTTTCGTCTTCCCGCGCTTCA |
| VPEAI_R | GACGAAAATAGCTTCGGGAACCCATTCCGCGGTGTACC |
| WFEEF_F | TGGTTTGAGGAGTTTTTCGTCTTCCCGCGCTTCA |
| WFEEF_R | GACGAAAAACTCCTCAAACCACCATTCCGCGGTGTACC |
| WMENI_F | TGGATGGAAAACATTTTCGTCTTCCCGCGCTTCA |
| WMENI_R | GACGAAAATGTTTTCCATCCACCATTCCGCGGTGTACC |
| EFERK_F | GAGTTTGAACGTAAATTCGTCTTCCCGCGCTTCA |
| EFERK_R | GACGAATTTACGTTCAAACTCCCATTCCGCGGTGTACC |
| KPEAD_F | AAGCCTGAGGCCGACTTCGTCTTCCCGCGCTTCA |
| KPEAD_R | GACGAAGTCGGCCTCAGGCTTCCATTCCGCGGTGTACC |
| ICEWN_F | ATTTGTGAATGGAACTTCGTCTTCCCGCGCTTCA |
| ICEWN_R | GACGAAGTTCCATTCACAAATCCATTCCGCGGTGTACC |
| CSERQ_F | TGTTCTGAACGTCAGTTCGTCTTCCCGCGCTTCA |
| CSERQ_R | GACGAACTGACGTTCAGAACACCATTCCGCGGTGTACC |
| HTEPQ_F | CACACCGAACCTCAGTTCGTCTTCCCGCGCTTCA |
| HTEPQ_R | GACGAACTGAGGTTCGGTGTGCCATTCCGCGGTGTACC |
| FGEWA_F | TTCGGAGAATGGGCTTTCGTCTTCCCGCGCTTCA |
| FGEWA_R | GACGAAAGCCCATTCTCCGAACCATTCCGCGGTGTACC |
| NVESN_F | AACGTAGAATCGAATTTCGTCTTCCCGCGCTTCA |
| NVESN_R | GACGAAATTCGATTCTACGTTCCATTCCGCGGTGTACC |
| MYEKL_F | ATGTATGAAAAGTTATTCGTCTTCCCGCGCTTCA |
| MYEKL_R | GACGAATAACTTTTCATACATCCATTCCGCGGTGTACC |
| SQEMQ_F | TCCCAGGAAATGCAGTTCGTCTTCCCGCGCTTCA |
| SQEMQ_R | GACGAACTGCATTTCCTGGGACCATTCCGCGGTGTACC |
| APEST_F | GCGCCGGAAAGCACTTTCGTCTTCCCGCGCTTCA |
| APEST_R | GACGAAAGTGCTTTCCGGCGCCCATTCCGCGGTGTACC |
| LMEQL_F | TTAATGGAGCAATTATTCGTCTTCCCGCGCTTCA |
| LMEQL_R | GACGAATAATTGCTCCATTAACCATTCCGCGGTGTACC |
| GTEIN_F | GGAACGGAAATTAACTTCGTCTTCCCGCGCTTCA |
| GTEIN_R | GACGAAGTTAATTTCCGTTCCCCATTCCGCGGTGTACC |
| WQEMV_F | TGGCAAGAGATGGTGTTCGTCTTCCCGCGCTTCA |
| WQEMV_R | GACGAACACCATCTCTTGCCACCATTCCGCGGTGTACC |
| GAEEA_F | GGGGCAGAAGAAGCTTTCGTCTTCCCGCGCTTCA |
| GAEEA_R | GACGAAAGCTTCTTCTGCCCCCCATTCCGCGGTGTACC |
| IIELF_F | ATTATCGAGTTATTCTTCGTCTTCCCGCGCTTCA |
| IIELF_R | GACGAAGAATAACTCGATAATCCATTCCGCGGTGTACC |

| | |
|---|---|
| AVESA_F | GCGGTCGAGAGCGCGTTCGTCTTCCCGCGCTTCA |
| AVESA_R | GACGAACGCGCTCTCGACCGCCCATTCCGCGGTGTACC |
| GIELG_F | GGTATTGAGCTGGGATTCGTCTTCCCGCGCTTCA |
| GIELG_R | GACGAATCCCAGCTCAATACCCCATTCCGCGGTGTACC |
| AEEVI_F | GCGGAGGAAGTTATCTTCGTCTTCCCGCGCTTCA |
| AEEVI_R | GACGAAGATAACTTCCTCCGCCCATTCCGCGGTGTACC |
| CLESI_F | TGTCTTGAAAGCATTTTCGTCTTCCCGCGCTTCA |
| CLESI_R | GACGAAAATGCTTTCAAGACACCATTCCGCGGTGTACC |
| DMEAV_F | GACATGGAGGCAGTCTTCGTCTTCCCGCGCTTCA |
| DMEAV_R | GACGAAGACTGCCTCCATGTCCCATTCCGCGGTGTACC |
| GWEVM_F | GGCTGGGAAGTCATGTTCGTCTTCCCGCGCTTCA |
| GWEVM_R | GACGAACATGACTTCCCAGCCCCATTCCGCGGTGTACC |
| AAEVF_F | GCCGCTGAGGTTTTCTTCGTCTTCCCGCGCTTCA |
| AAEVF_R | GACGAAGAAAACCTCAGCGGCCCATTCCGCGGTGTACC |
| GNEVI_F | GGCAACGAAGTGATTTTCGTCTTCCCGCGCTTCA |
| GNEVI_R | GACGAAAATCACTTCGTTGCCCCATTCCGCGGTGTACC |
| ALELV_F | GCCCTTGAGTTAGTGTTCGTCTTCCCGCGCTTCA |
| ALELV_R | GACGAACACTAACTCAAGGGCCCATTCCGCGGTGTACC |
| **Library 3 (WYTAEWGLEXXXXXXRFI)** | |
| FKPKEE_F | TTCAAGCCGAAGGAGGAGCGCTTCATCTGAGCGGC |
| FKPKEE_R | CTCCTCCTTCGGCTTGAACTCGAGGCCCCATTCC |
| KQCNMM_F | AAACAGTGTAATATGATGCGCTTCATCTGAGCGGC |
| KQCNMM_R | CATCATATTACACTGTTTCTCGAGGCCCCATTCC |
| GWRNTT_F | GGATGGCGCAATACGACGCGCTTCATCTGAGCGGC |
| GWRNTT_R | CGTCGTATTGCGCCATCCCTCGAGGCCCCATTCC |
| NWRMEI_F | AATTGGCGCATGGAGATTCGCTTCATCTGAGCGGC |
| NWRMEI_R | AATCTCCATGCGCCAATTCTCGAGGCCCCATTCC |
| TLEFPR_F | ACCCTTGAGTTCCCACGCCGCTTCATCTGAGCGGC |
| TLEFPR_R | GCGTGGGAACTCAAGGGTCTCGAGGCCCCATTCC |
| LYGTCD_F | CTTTACGGGACATGCGATCGCTTCATCTGAGCGGC |
| LYGTCD_R | ATCGCATGTCCCGTAAAGCTCGAGGCCCCATTCC |
| WGNTFN_F | TGGGGAAATACTTTTAATCGCTTCATCTGAGCGGC |
| WGNTFN_R | ATTAAAAGTATTTCCCCACTCGAGGCCCCATTCC |
| ECIQAR_F | GAATGCATCCAGGCCCGTCGCTTCATCTGAGCGGC |
| ECIQAR_R | ACGGGCCTGGATGCATTCCTCGAGGCCCCATTCC |
| RWMMMG_F | CGCTGGATGATGATGGGACGCTTCATCTGAGCGGC |
| RWMMMG_R | TCCCATCATCATCCAGCGCTCGAGGCCCCATTCC |
| TNPPCF_F | ACAAACCCACCTTGTTTCCGCTTCATCTGAGCGGC |
| TNPPCF_R | GAAACAAGGTGGGTTTGTCTCGAGGCCCCATTCC |
| PLFMPT_F | CCCTTGTTTATGCCCACACGCTTCATCTGAGCGGC |

| | |
|---|---|
| PLFMPT_R | TGTGGGCATAAACAAGGGCTCGAGGCCCCATTCC |
| TKCCAY_F | ACCAAGTGCTGTGCTTATCGCTTCATCTGAGCGGC |
| TKCCAY_R | ATAAGCACAGCACTTGGTCTCGAGGCCCCATTCC |
| TYGVGG_F | ACCTACGGAGTTGGTGGACGCTTCATCTGAGCGGC |
| TYGVGG_R | TCCACCAACTCCGTAGGTCTCGAGGCCCCATTCC |
| SLHPYI_F | TCGCTGCATCCCTACATTCGCTTCATCTGAGCGGC |
| SLHPYI_R | AATGTAGGGATGCAGCGACTCGAGGCCCCATTCC |
| YLFFCQ_F | TACTTATTTTTTTGTCAGCGCTTCATCTGAGCGGC |
| YLFFCQ_R | CTGACAAAAAAATAAGTACTCGAGGCCCCATTCC |
| QTAMDY_F | CAAACAGCGATGGATTACCGCTTCATCTGAGCGGC |
| QTAMDY_R | GTAATCCATCGCTGTTTGCTCGAGGCCCCATTCC |
| AFVLHA_F | GCTTTCGTCCTTCATGCTCGCTTCATCTGAGCGGC |
| AFVLHA_R | AGCATGAAGGACGAAAGCCTCGAGGCCCCATTCC |
| GTKWFT_F | GGGACCAAGTGGTTTACGCGCTTCATCTGAGCGGC |
| GTKWFT_R | CGTAAACCACTTGGTCCCCTCGAGGCCCCATTCC |
| LWFCSP_F | TTGTGGTTTTGCAGCCCGCGCTTCATCTGAGCGGC |
| LWFCSP_R | CGGGCTGCAAAACCACAACTCGAGGCCCCATTCC |
| LITLLN_F | TTGATTACCTTGCTTAACCGCTTCATCTGAGCGGC |
| LITLLN_R | GTTAAGCAAGGTAATCAACTCGAGGCCCCATTCC |
| LNFFCP_F | TTGAATTTCTTCTGTCCGCGCTTCATCTGAGCGGC |
| LNFFCP_R | CGGACAGAAGAAATTCAACTCGAGGCCCCATTCC |
| YLFYMT_F | TACCTTTTCTACATGACACGCTTCATCTGAGCGGC |
| YLFYMT_R | TGTCATGTAGAAAAGGTACTCGAGGCCCCATTCC |
| IFWGIN_F | ATTTTTTGGGGTATTAATCGCTTCATCTGAGCGGC |
| IFWGIN_R | ATTAATACCCCAAAAAATCTCGAGGCCCCATTCC |
| MYWNVP_F | ATGTACTGGAATGTGCCACGCTTCATCTGAGCGGC |
| MYWNVP_R | TGGCACATTCCAGTACATCTCGAGGCCCCATTCC |
| VFGLLT_F | GTGTTTGGTTTGTTGACGCGCTTCATCTGAGCGGC |
| VFGLLT_R | CGTCAACAAACCAAACACCTCGAGGCCCCATTCC |
| MWDVLP_F | ATGTGGGACGTACTTCCACGCTTCATCTGAGCGGC |
| MWDVLP_R | TGGAAGTACGTCCCACATCTCGAGGCCCCATTCC |
| MGVAFC_F | ATGGGTGTCGCATTCTGTCGCTTCATCTGAGCGGC |
| MGVAFC_R | ACAGAATGCGACACCCATCTCGAGGCCCCATTCC |
| MFLFMS_F | ATGTTCCTGTTCATGTCCCGCTTCATCTGAGCGGC |
| MFLFMS_R | GGACATGAACAGGAACATCTCGAGGCCCCATTCC |
| TAMAFG_F | ACGGCCATGGCATTTGGCCGCTTCATCTGAGCGGC |
| TAMAFG_R | GCCAAATGCCATGGCCGTCTCGAGGCCCCATTCC |
| TLMYAP_F | ACCTTGATGTACGCACCTCGCTTCATCTGAGCGGC |
| TLMYAP_R | AGGTGCGTACATCAAGGTCTCGAGGCCCCATTCC |
| **Chimeric peptide primers** | |

| | |
|---|---|
| pET28_gib_F | ACTGAGATCCGGCTGCTAAC |
| pET28_gib_R | GGATCCGGATTGGAAGTACAG |
| FusALP_gib_F | GAACCTGTACTTCCAATCCGGATCCATGGTACGCC |
| BulA_gib_R | CTTTGTTAGCAGCCGGATCTCAGTTTAATGAAAACGCACTCCAGTC |
| EndA_gib_R | CTTTGTTAGCAGCCGGATCTCAGTTTAAATCATGCGTGGCAGG |
| Cau31A_gib_F | CTTTGTTAGCAGCCGGATCTCAGTTTAAGCGAAATAGTATTGGCTAAC |
| FlaA_gib_R | CTTTGTTAGCAGCCGGATCTCAGTTTACCCTGGATCATAGTGAAAGG |
| RegA_gib_R | CTTTGTTAGCAGCCGGATCTCAGTTTATAACGAGAAGCGAGCCG |
| Sma6A_gib_R | CTTTGTTAGCAGCCGGATCTCAGTTTAATCATTAGAGATTCCCACCTG |
| MflA_gib_R | CTTTGTTAGCAGCCGGATCTCAGTTTAGTTGATACGGCTGTTAGG |
| XylA_gib_R | CTTTGTTAGCAGCCGGATCTCAGTTTAAATCCAGGTATCGAAGAACC |
| SegA_gib_R | CTTTGTTAGCAGCCGGATCTCAGTTTATTCTTGTGTTAATCCAGCTGG |
| ChloA_gib_R | CTTTGTTAGCAGCCGGATCTCAGTTTACCCCCCAAAACGC |
| EnsA_gib_R | CTTTGTTAGCAGCCGGATCTCAGTTTAAAATGTGCGGTAACGCTTC |
| SidA_gib_R | CTTTGTTAGCAGCCGGATCTCAGTTTACACGATACGGCGTAC |
| RhoA_gib_R | CTTTGTTAGCAGCCGGATCTCAGTTTAATTGAATTTTCCGACCCAC |
| CarA_gib_R | CTTTGTTAGCAGCCGGATCTCAGTTTATACAATGCGACGGGAC |
| CacA_gib_R | CTTTGTTAGCAGCCGGATCTCAGTTTACTGGTCACGGCTC |
| KorA_gib_R | CTTTGTTAGCAGCCGGATCTCAGTTTATGGTTTACGCATCATCATCG |
| OliA_gib_R | CTTTGTTAGCAGCCGGATCTCAGTTTATGAATCACGTTCCAAGATGAATTC |
| BacA_gib_R | CTTTGTTAGCAGCCGGATCTCAGTTTACACTTTCCAATCCATAAGAATC |
| HaiA_gib_R | CTTTGTTAGCAGCCGGATCTCAGTTTATGTGATTTTGTACCCGTACAAG |
| KunA_gib_R | CTTTGTTAGCAGCCGGATCTCAGTTTAAGGTTTGAAGTACAACTGCTC |
| NetA_gib_R | CTTTGTTAGCAGCCGGATCTCAGTTTACATGCGCTTCAACAGC |
| JesA_gib_R | CTTTGTTAGCAGCCGGATCTCAGTTTAACTGGGCGTGAATAACG |
| PpcA_gib_R | CTTTGTTAGCAGCCGGATCTCAGTTTATACGCGGTATGCGTAAATTAAG |
| SulA_gib_R | CTTTGTTAGCAGCCGGATCTCAGTTCAGGAATTATGG |
| BbaA_gib_R | CTTTGTTAGCAGCCGGATCTCAGTTCAGCCTTCTAC |
| SalA_gib_R | CTTTGTTAGCAGCCGGATCTCAGTTCAGAACCAGCTATG |
| **Chimeric peptide ultramers** | |
| BulA | GGATCCATGGAAAAGAAGAAGTACACTGCACCTCAATTGGCCAAGGTGGGCGAGT TTAAGGAAGCGACTGGGGCTGATTCGCCAGATGGCGAGTTTGAACTTATTCCAGT ACAGCGCTGGACAATGACTGGAGTGCGTTTTCATTAAAAGCTT |
| EndA | GGATCCATGGAAAAGAAGAAGTACACTGCACCTCAATTGGCCAAGGTGGGCGAGT TTAAGGAAGCGACTGGGTGGCAGCAAGGGCGTGGATTTGAGGTTCTTTTTTTCCT GCCACGCATGATTTAAAAGCTT |
| Cau31A | GGATCCATGGAAAAGAAGAAGTACACTGCACCTCAATTGGCCAAGGTGGGCGAGT TTAAGGAAGCGACTGGGTCCTTTGACGTGGGGACTATTAAAGAAGGTTTAGTTAG CCAATACTATTTCGCTTAAAAGCTT |

| | |
|---|---|
| FlaA | GGATCCATGGAAAAGAAGAAGTACACTGCACCTCAATTGGCCAAGGTGGGCGAGT<br>TTAAGGAAGCGACTGGGGGTGGTGATGGCCCCGGCACGGAGATGCTTACCTTTCA<br>CTATGATCCAGGGTAAAAGCTT |
| RegA | GGATCCATGGAAAAGAAGAAGTACACTGCACCTCAATTGGCCAAGGTGGGCGAGT<br>TTAAGGAAGCGACTGGGCTTGGCACCCACCGTGGTCCTGAGCGTGTTTTACCGGC<br>TCGCTTCTCGTTATAAAAGCTT |
| Sma6A | GGATCCATGGAAAAGAAGAAGTACACTGCACCTCAATTGGCCAAGGTGGGCGAGT<br>TTAAGGAAGCGACTGGGGGTTCCATTGGAGCACTTGGCGATGAAAACGGATTGCA<br>CAAACAGGTGGGAATCTCTAATGATTAAAAGCTT |
| MflA | GGATCCATGGAAAAGAAGAAGTACACTGCACCTCAATTGGCCAAGGTGGGCGAGT<br>TTAAGGAAGCGACTGGGGGTGATATTTATCCGGGGGTTGAAAGTCTGGATCCTAA<br>CAGCCGTATCAACTAAAAGCTT |
| XylA | GGATCCATGGAAAAGAAGAAGTACACTGCACCTCAATTGGCCAAGGTGGGCGAGT<br>TTAAGGAAGCGACTGGGGTGTTTTATGTTCGCAACGGCGAAGAGGTACTTTGGTT<br>CTTCGATACCTGGATTTAAAAGCTT |
| SegA | GGATCCATGGAAAAGAAGAAGTACACTGCACCTCAATTGGCCAAGGTGGGCGAGT<br>TTAAGGAAGCGACTGGGGGTATGCCTGGCGCCTTTGTGGAGATTCTGGGCGAGGA<br>TGATAAGCCAGCTGGATTAACACAAGAATAAAAGCTT |
| ChloA | GGATCCATGGAAAAGAAGAAGTACACTGCACCTCAATTGGCCAAGGTGGGCGAGT<br>TTAAGGAAGCGACTGGGGCCTCCATGAACGAGATTGCACCGGAACTTGTAGGCGA<br>CAAGACACAGCGTTTTGGGGGGTAAAAGCTT |
| EnsA | GGATCCATGGAAAAGAAGAAGTACACTGCACCTCAATTGGCCAAGGTGGGCGAGT<br>TTAAGGAAGCGACTGGGGGGAACAATAGTGCAGGCGTCCATGAGACCTTGGGCCC<br>GAAGCGTTACCGCACATTTTAAAAGCTT |
| SidA | GGATCCATGGAAAAGAAGAAGTACACTGCACCTCAATTGGCCAAGGTGGGCGAGT<br>TTAAGGAAGCGACTGGGGGGTACTTTGTTGGAAGCTATAAGGAGTGGATTGTACG<br>CCGTATCGTGTAAAAGCTT |
| RhoA | GGATCCATGGAAAAGAAGAAGTACACTGCACCTCAATTGGCCAAGGTGGGCGAGT<br>TTAAGGAAGCGACTGGGGGCGGAGGTATTTGGTGGGTCGAGTGGGTCGGAAAATT<br>CAATTAAAAGCTT |
| CarA | GGATCCATGGAAAAGAAGAAGTACACTGCACCTCAATTGGCCAAGGTGGGCGAGT<br>TTAAGGAAGCGACTGGGGGATACTTCTATGGCAGTTATAAGGAATTCTTGTCCCG<br>TCGCATTGTATAAAAGCTT |
| CacA | GGATCCATGGAAAAGAAGAAGTACACTGCACCTCAATTGGCCAAGGTGGGCGAGT<br>TTAAGGAAGCGACTGGGGGATATCCCCTTGGCGCTCAAGAAATCGTTGGGTTCTT<br>GAGCCGTGACCAGTAAAAGCTT |
| KorA | GGATCCATGGAAAAGAAGAAGTACACTGCACCTCAATTGGCCAAGGTGGGCGAGT<br>TTAAGGAAGCGACTGGGTCAGGTAGTCTGACCCCCACAGAATCAATGACGATGAT<br>GATGCGTAAACCATAAAAGCTT |
| OliA | GGATCCATGGAAAAGAAGAAGTACACTGCACCTCAATTGGCCAAGGTGGGCGAGT<br>TTAAGGAAGCGACTGGGGGTGAAATGGGGACGTTGCTGGAATTCATCTTGGAACG<br>TGATTCATAAAAGCTT |

| | |
|---|---|
| BacA | GGATCCATGGAAAAGAAGAAGTACACTGCACCTCAATTGGCCAAGGTGGGCGAGT<br>TTAAGGAAGCGACTGGGGCAGGTAGTAAGGGATATCAAGAAGTGGTCATGATTCT<br>TATGGATTGGAAAGTGTAAAAGCTT |
| HaiA | GGATCCATGGAAAAGAAGAAGTACACTGCACCTCAATTGGCCAAGGTGGGCGAGT<br>TTAAGGAAGCGACTGGGGGCCTGCCCTGGACCCGTACCGAAGCCTTGTACGGGTA<br>CAAAATCACATAAAAGCTT |
| KunA | GGATCCATGGAAAAGAAGAAGTACACTGCACCTCAATTGGCCAAGGTGGGCGAGT<br>TTAAGGAAGCGACTGGGGGGAACCCAGATGGTGACGAAGTTGAGATCGACGAGCA<br>GTTGTACTTCAAACCTTAAAAGCTT |
| NetA | GGATCCATGGAAAAGAAGAAGTACACTGCACCTCAATTGGCCAAGGTGGGCGAGT<br>TTAAGGAAGCGACTGGGTTGGGCATTTCAGGGGGACCAGAACTGCTGCTGTTGAA<br>GCGCATGTAAAAGCTT |
| JesA | GGATCCATGGAAAAGAAGAAGTACACTGCACCTCAATTGGCCAAGGTGGGCGAGT<br>TTAAGGAAGCGACTGGGACAGGCCCGAAGACTTTCACTCAAGAAATCTTGACGTT<br>ATTCACGCCCAGTTAAAAGCTT |
| PpcA | GGATCCATGGAAAAGAAGAAGTACACTGCACCTCAATTGGCCAAGGTGGGCGAGT<br>TTAAGGAAGCGACTGGGGCCCCTGCGATGTTCGGTTCTCCAGAAATCGGGAACTT<br>AATTTACGCATACCGCGTATAAAAGCTT |
| BbaA | GGATCCATGGAAAAGAAGAAGTACACTGCACCTCAATTGGCCAAGGTGGGCGAGT<br>TTAAGGAAGCGACTGGGGATGCCTTACCCGGTCCCTATTTAGAAATGGGGATATT<br>CCCGAGCAGAACAGTAGAAGGCTGAAAGCTT |
| SalA | GGATCCATGGAAAAGAAGAAGTACACTGCACCTCAATTGGCCAAGGTGGGCGAGT<br>TTAAGGAAGCGACTGGGTTCATTGGTCCTATCCATTACGAAGGTATATTGTTATG<br>GCATAGCTGGTTCTGAAAGCTT |
| SulA | GGATCCATGGAAAAGAAGAAGTACACTGCACCTCAATTGGCCAAGGTGGGCGAGT<br>TTAAGGAAGCGACTGGGAGCGGTGCCCTTCTGGACTTGGTAGAAGTATTCCTTGT<br>CTCCCATAATTCCTGAAAGCTT |

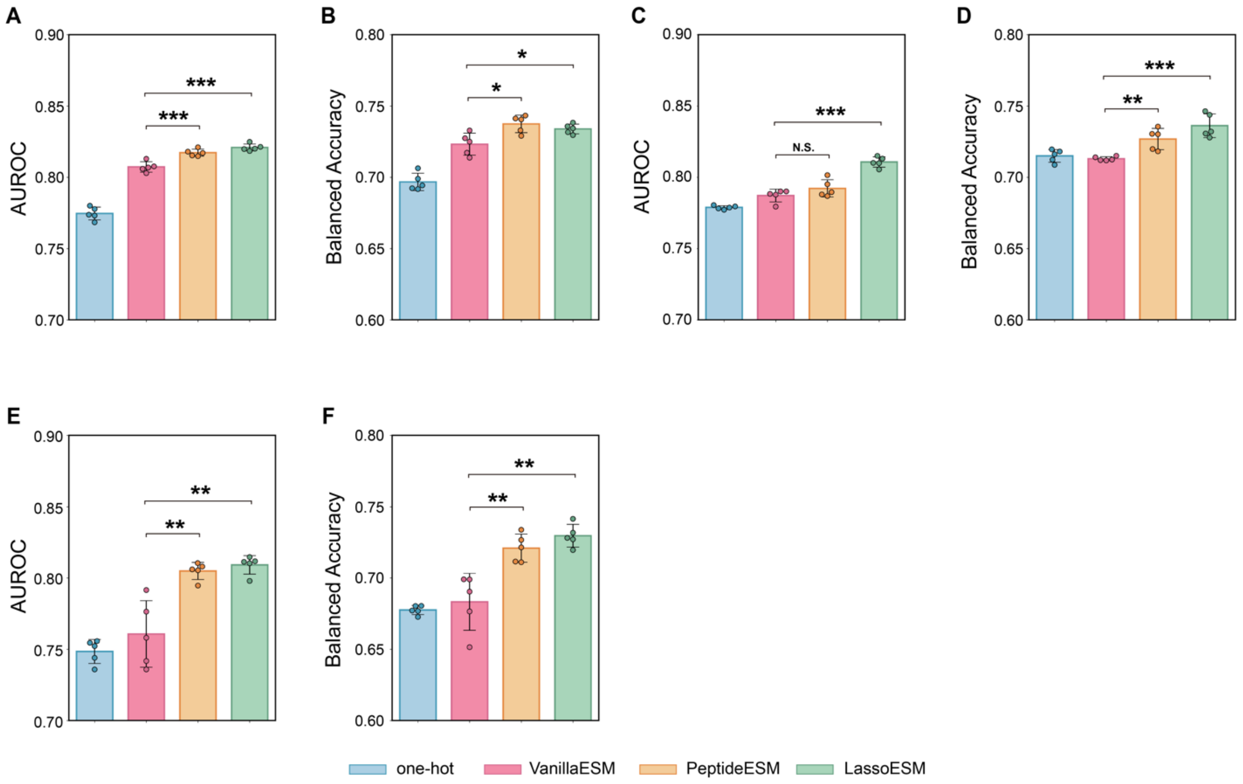


**Figure S1**. **Evaluation of different classification models trained on different embeddings of the fusilassin variants dataset.** AUROC score and balanced accuracy of Random Forest **(A, B)**, Adaptive Boosting **(C, D)**, Multilayer Perceptron **(E, F)** for the fusilassin dataset trained on embeddings from one-hot encoding (blue), VanillaESM[1] (pink), PeptideESM[2] (orange) and LassoESM (green). Error band depicts mean ± standard deviation calculated over 5 replicates of 10-fold cross validation. $p$-value was calculated using two-sided t-test. *$p < 0.05$, ** $p < 0.01$, *** $p < 0.001$.

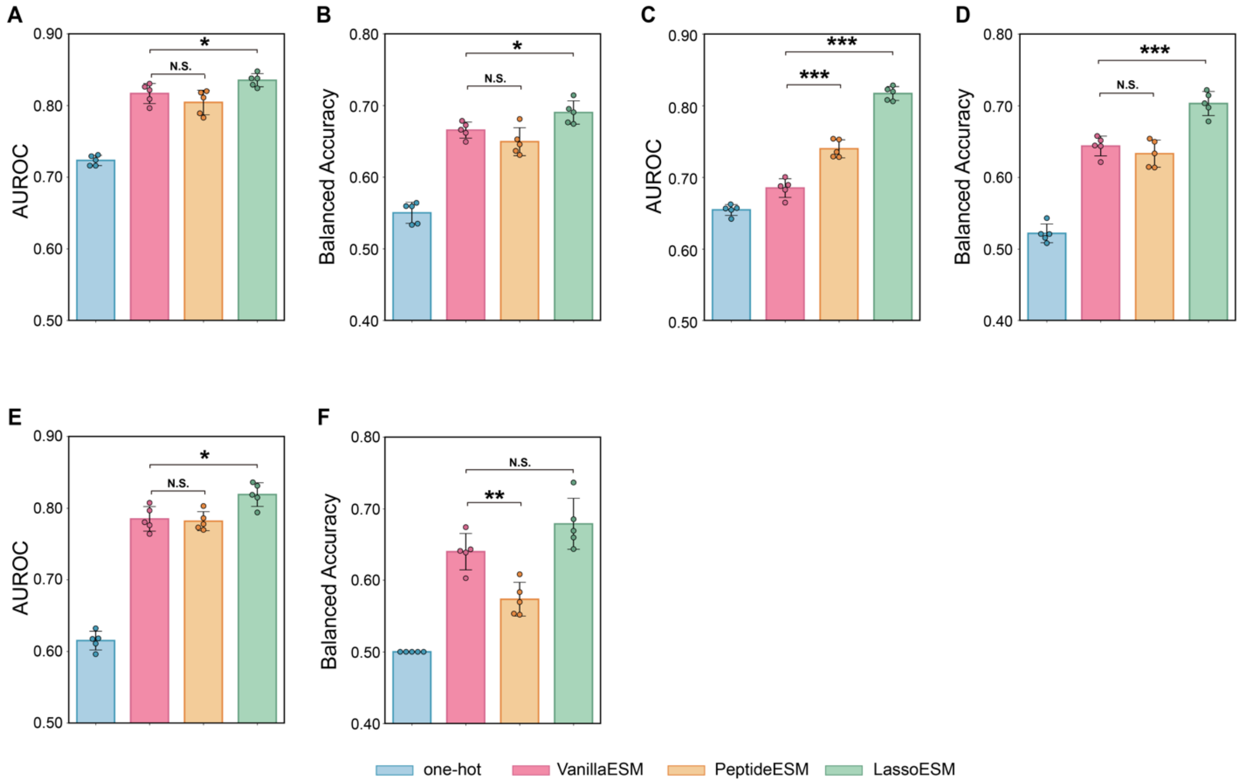


**Figure S2**. **Evaluation of different classification models trained on different embeddings of the microcinJ25 variants dataset.** AUROC score and balanced accuracy of Random Forest **(A, B)**, Adaptive Boosting **(C, D)**, Multilayer Perceptron **(E, F)** for the microcin J25 dataset trained on embeddings from one-hot encoding (blue), VanillaESM[1] (pink), PeptideESM[2] (orange) and LassoESM (green). Error band depicts mean ± standard deviation calculated over 5 repeats of 10-fold cross validation. *p*-value was calculated using two-sided t-test. *$p < 0.05$, ** $p < 0.01$, *** $p < 0.001$.

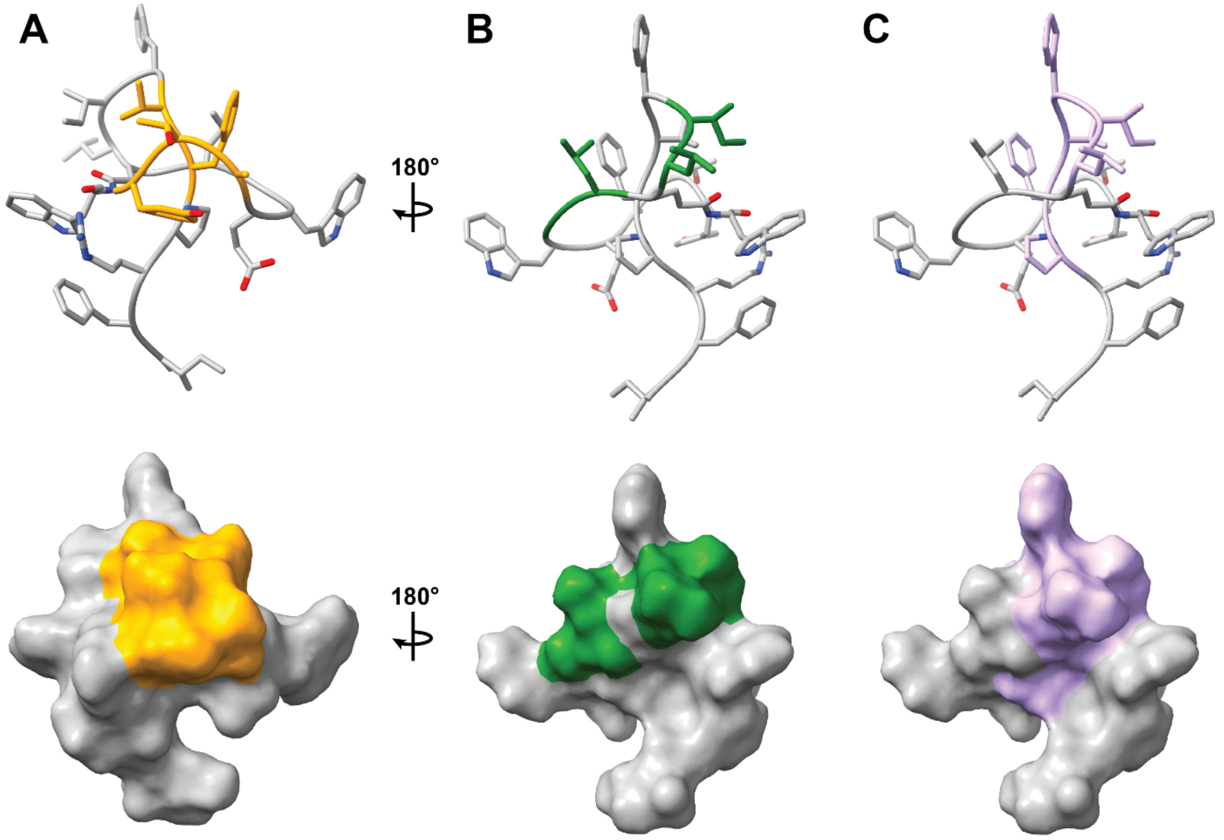


**Figure S3. Fusilassin model in stick and surface representation.** Figure shows varied positions in library 1 (A), library 2 (B), and library 3 (C). The model was generated using LassoHTP.[4]

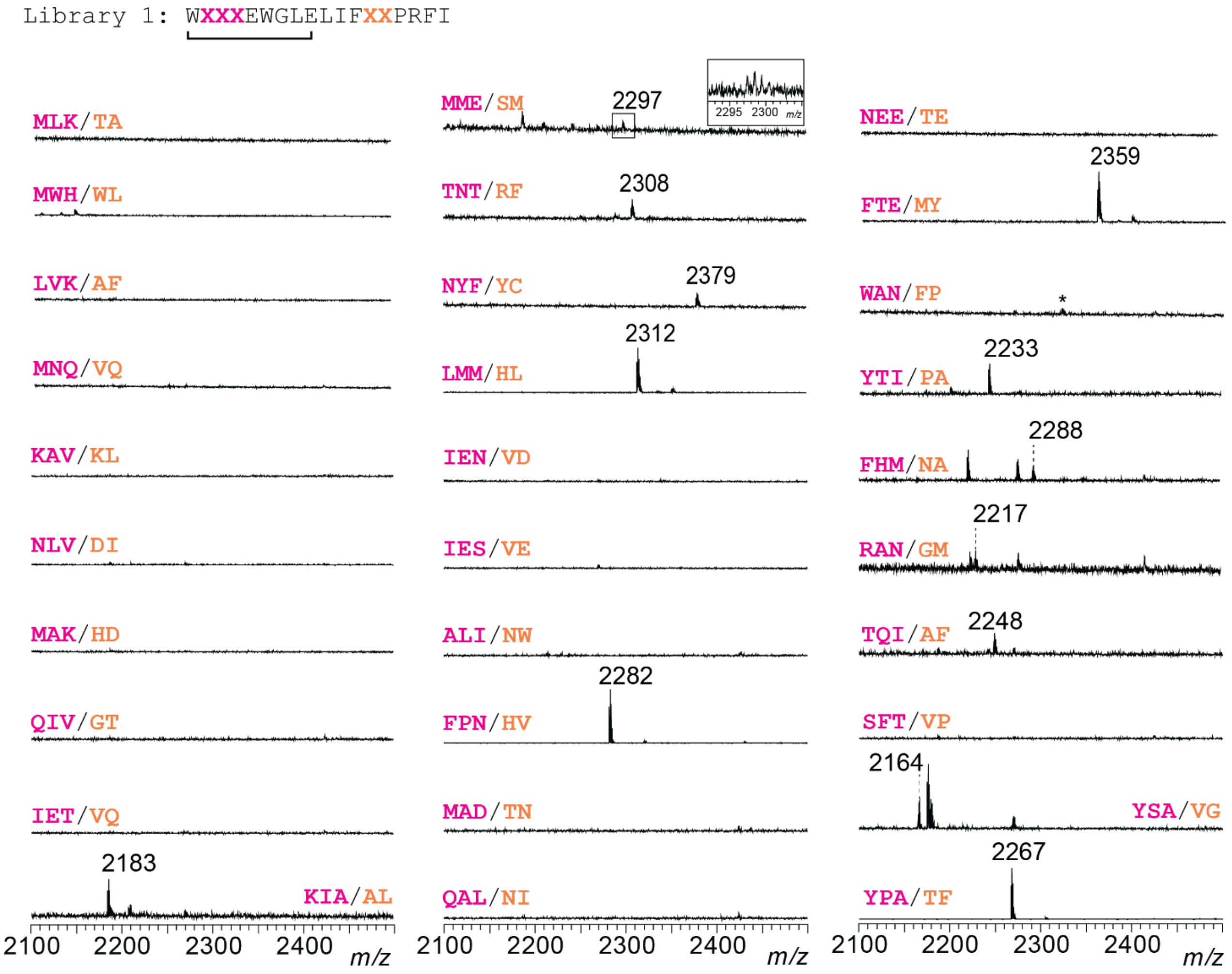


**Figure S4. MALDI-TOF-MS analysis of randomly selected round 1 fusilassin variants from library 1 tested using CFB.** Library 1 contains variations at positions 2, 3, and 4 (pink), and positions 13 and 14 (orange). A numeric value is given for the $[M+H]^+$ of the cyclized lasso peptide when observed. In cases where the spectrum contains several peaks in the mass window, a dashed line indicates the peak corresponding to the cyclized mass. * indicates the $[M+H]^+$ for the linear core peptide. Spectra correspond to sequences in **Table S4**.

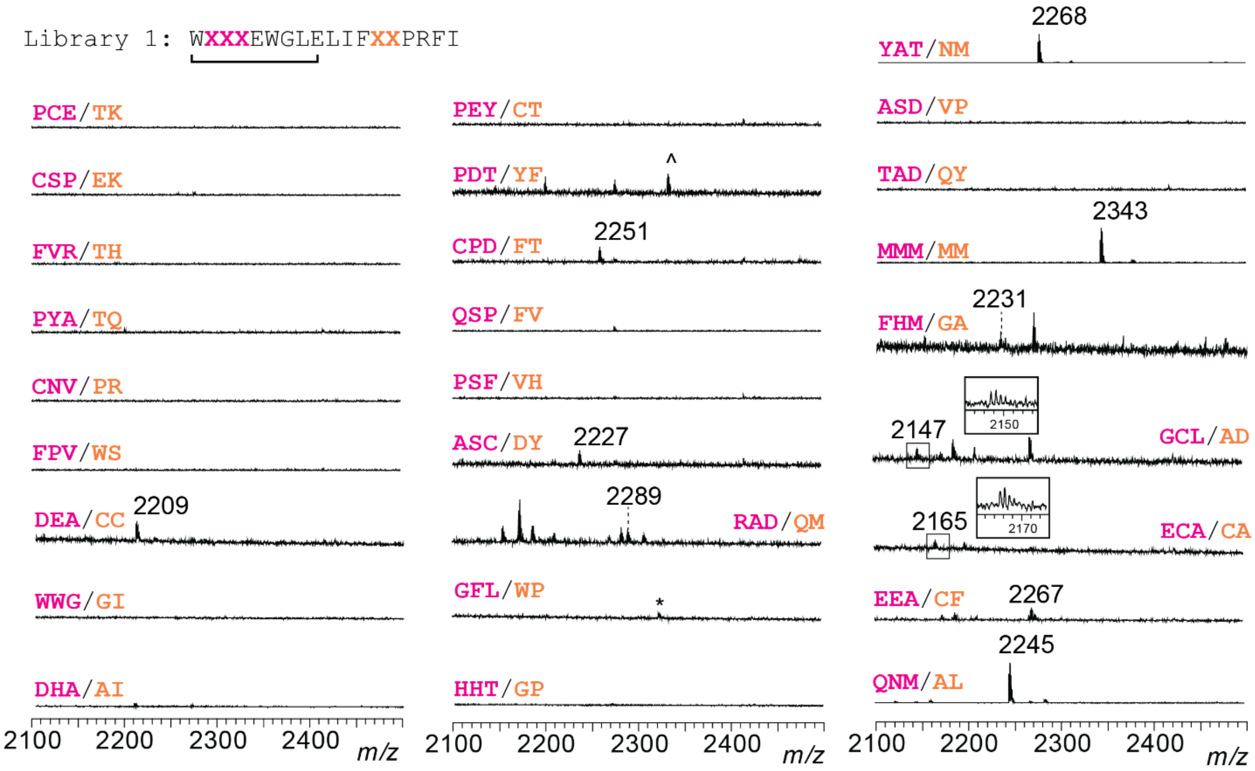


**Figure S5. MALDI-TOF-MS analysis of round 1 fusilassin variants for library 1 tested using CFB**. Positions 2, 3, and 4 are colored pink, while positions 13 and 14 are colored orange. The $[M+H]^+$ for the cyclized lasso peptide is labeled. In cases where the spectrum contains several peaks in the mass window, a dashed line indicates the peak corresponding to the cyclized mass. * indicates the $[M+H]^+$ for the linear core peptide. ^ indicates a $[M+Na]^+$ peak for the cyclized lasso peptide. Insets with zoomed-in spectra are provided for small peaks. Spectra correspond to sequences on **Table S5**.

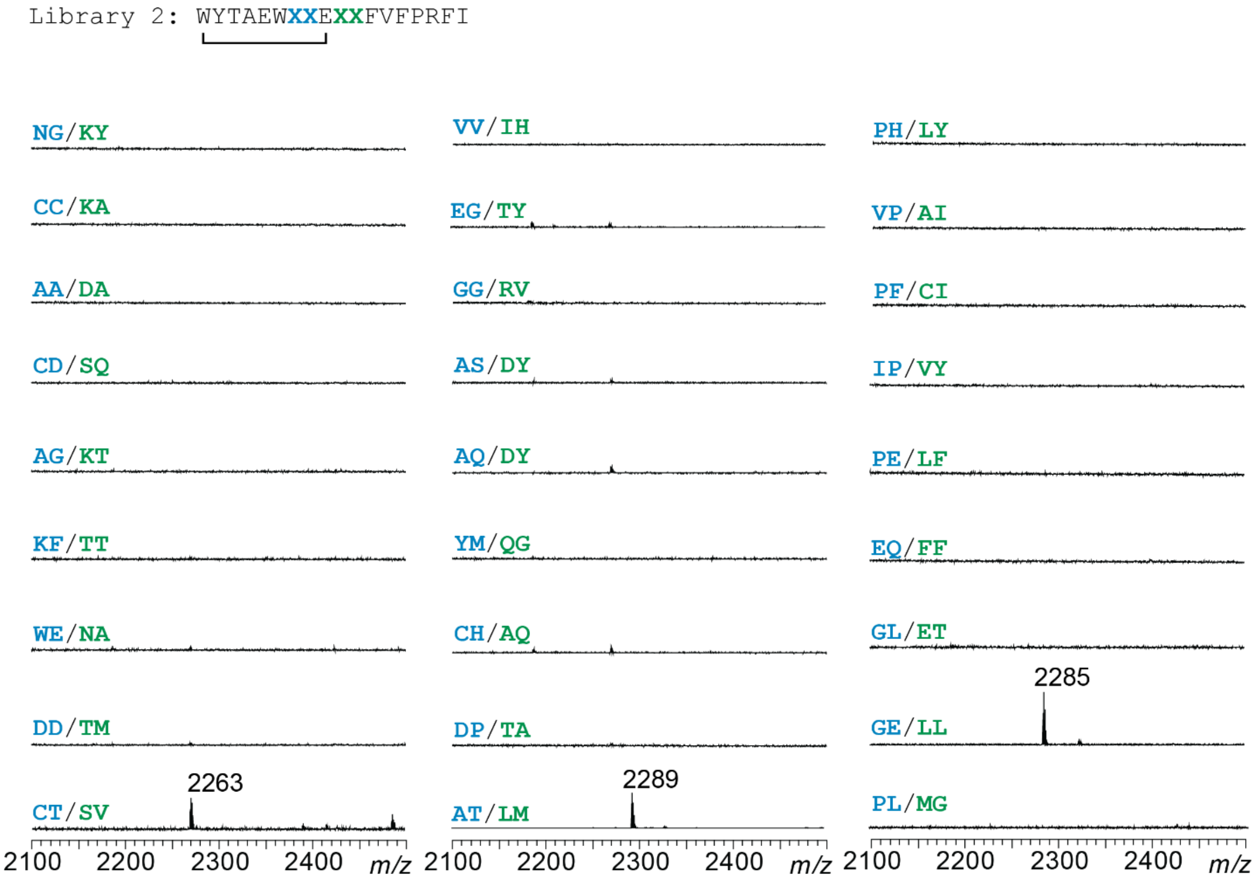


**Figure S6. MALDI-TOF-MS analysis of round 1 fusilassin variants from library 2 tested using CFB.** Library 2 contains variation at positions 7 and 8, which are colored blue, and positions 10 and 11, which are colored green. The $[M+H]^+$ for the cyclized lasso peptide is labeled. Spectra correspond to sequences on **Table S6**.

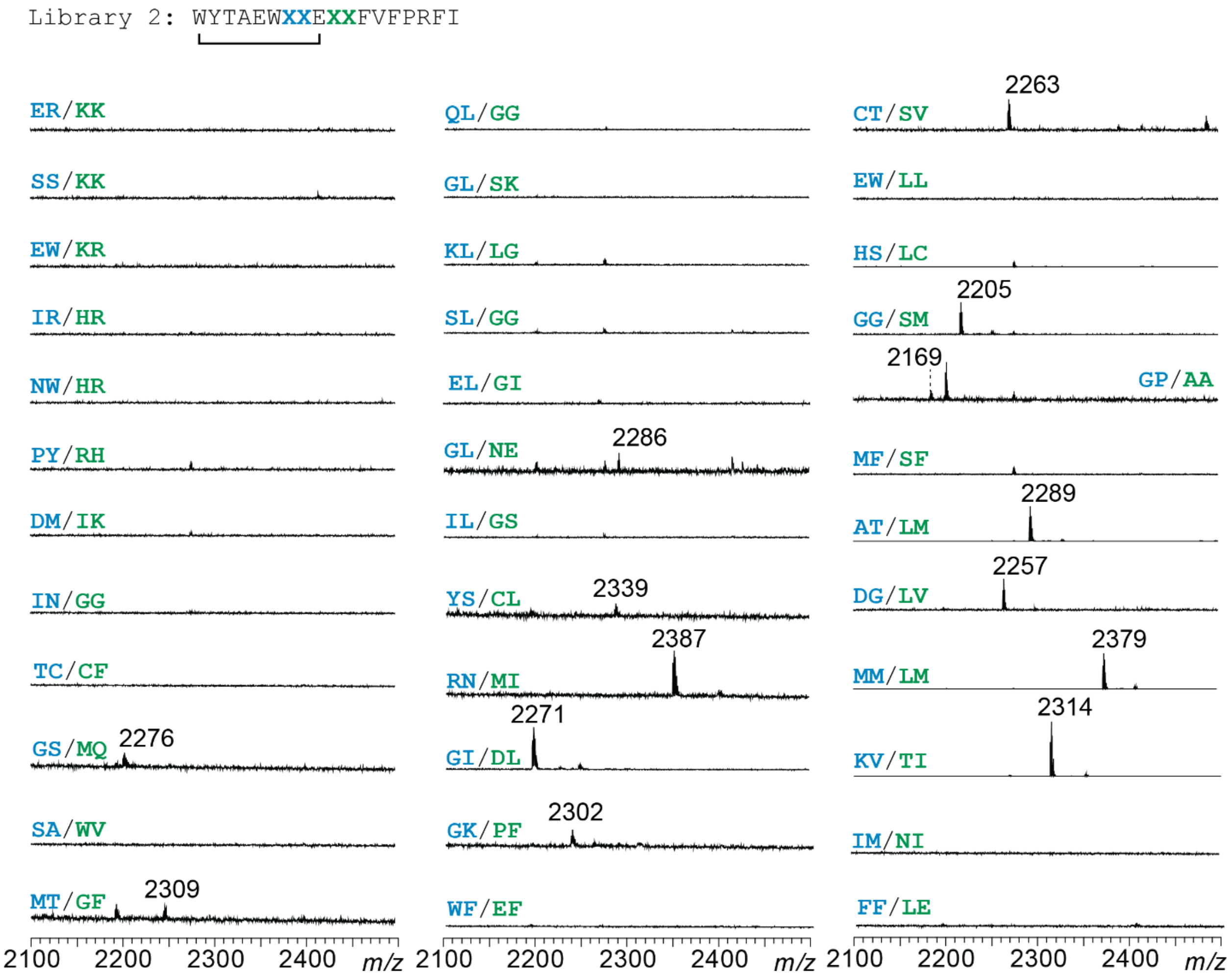


**Figure S7. MALDI-TOF-MS analysis of round 1 fusilassin variants for library 2 tested using CFB.** Library 2 contains variation at positions 7 and 8, which are colored blue, and positions 10 and 11, which are colored green. The $[M+H]^+$ for the cyclized lasso peptide is labeled. In cases where the spectrum contains several peaks in the mass window, a dashed line indicates the peak corresponding to the cyclized mass. Spectra correspond to sequences on **Table S7**.

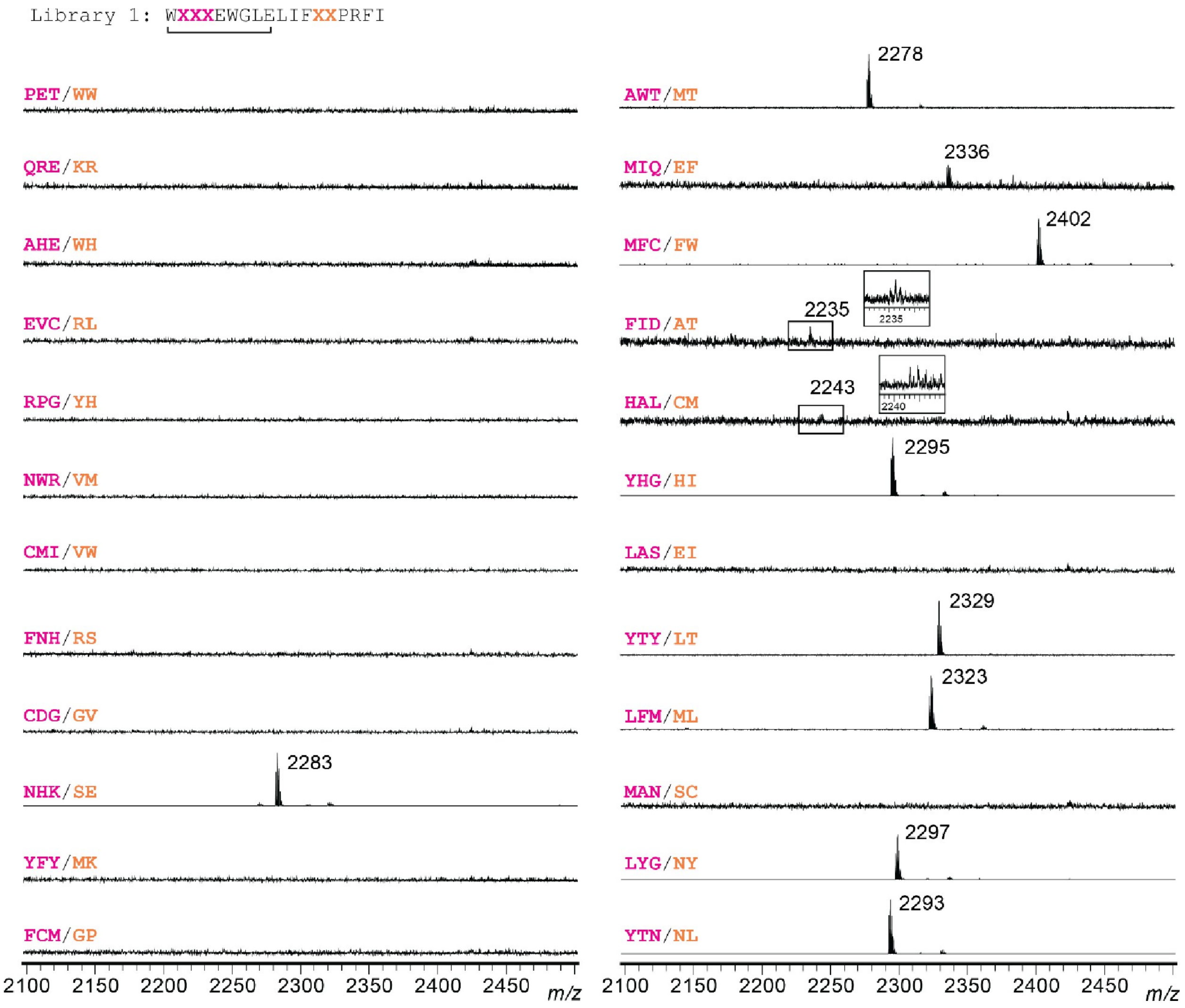


**Figure S8**. **MALDI-TOF-MS analysis of round 2 fusilassin variants for library 1 tested using CFB.** Positions 2, 3, and 4 are colored pink, while positions 13 and 14 are colored orange. The $[M+H]^+$ for the cyclized lasso peptide is labeled. Insets with zoomed-in spectra are provided for small peaks. Spectra correspond to sequences on **Table S8**.

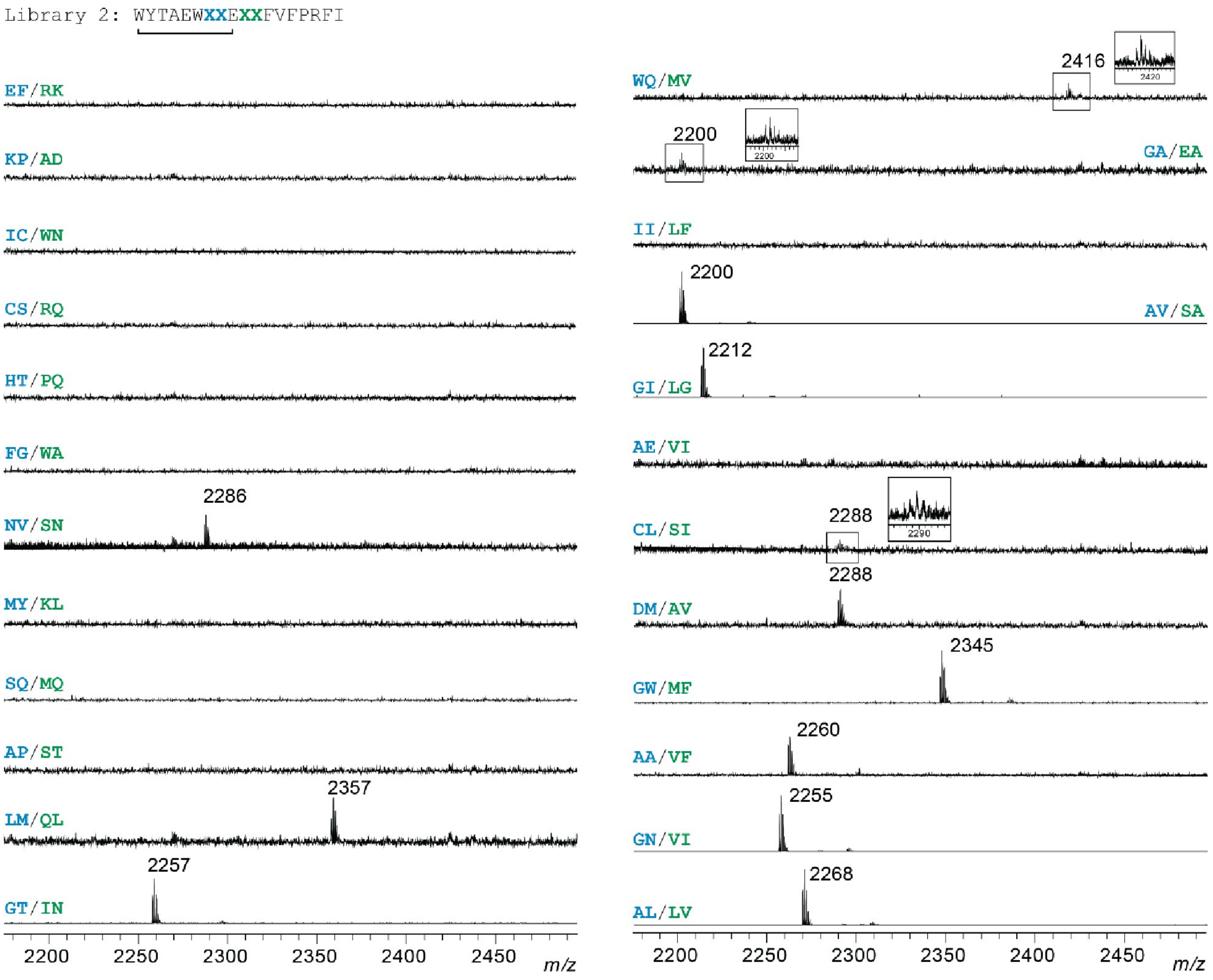


**Figure S9. MALDI-TOF-MS analysis of round 2 fusilassin variants for library 2 tested using CFB.** Positions 7 and 8 are blue, while positions 10 and 11 are green. The $[M+H]^+$ for the cyclized lasso peptide is labeled. Insets with zoomed-in spectra are provided for small peaks. Spectra correspond to sequences on **Table S9**.

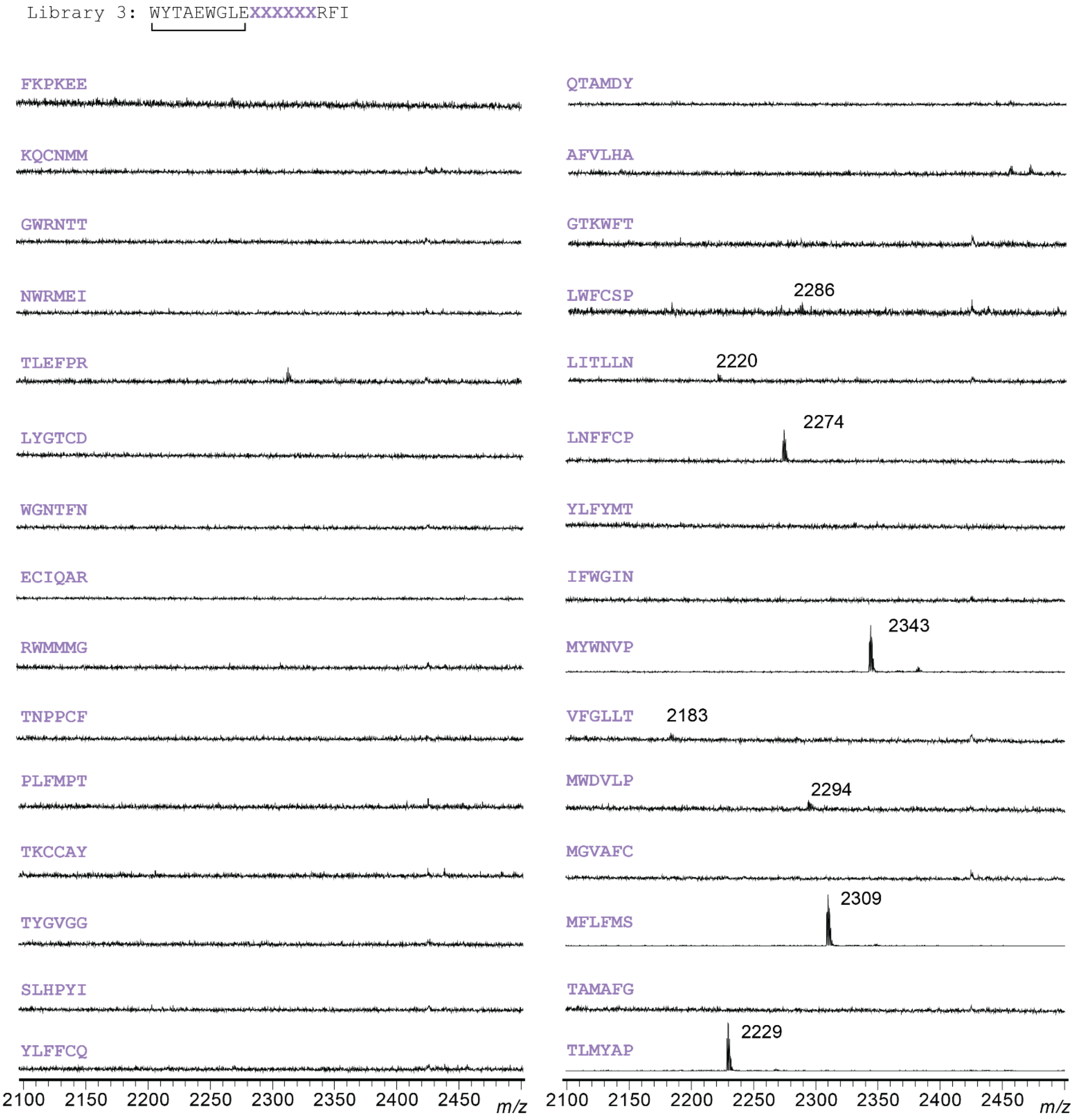


**Figure S10. MALDI-TOF-MS analysis of fusilassin variants for library 3 tested using CFB.** Positions 10-15 are purple. The $[M+H]^+$ for the cyclized lasso peptide is labeled. Spectra correspond to sequences on **Table S10**.

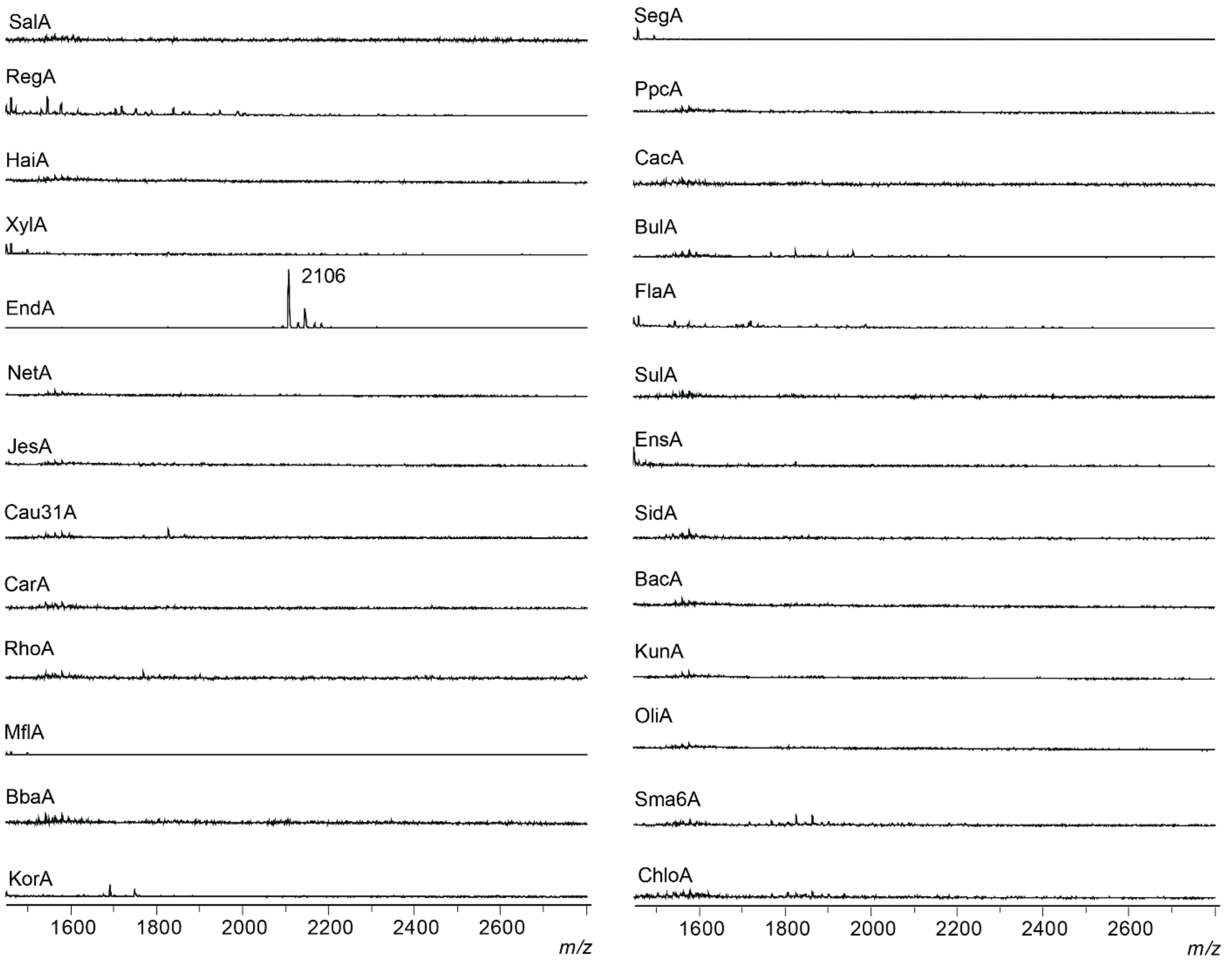


**Figure S11. MALDI-TOF-MS analysis of randomly selected chimeric core peptides tested using CFB.** Precursor peptides contain the FusA leader peptide and a chimeric core peptide. All sequences were tested with the fusilassin cyclase to examine its compatibility with chimeric core peptide sequences. Spectra correspond to sequences shown in **Table S11**.

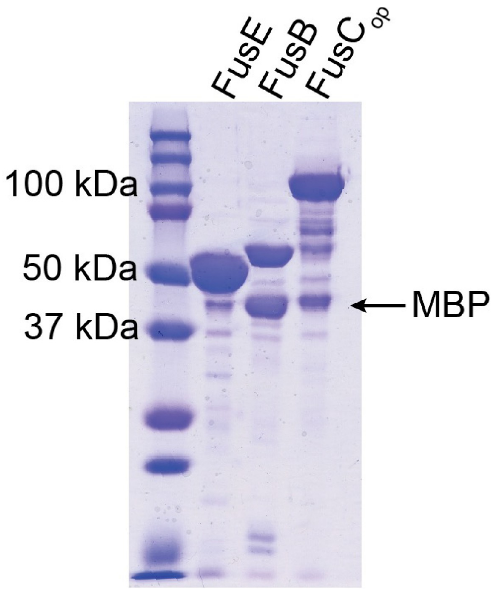


**Figure S12. SDS-PAGE gel for purified proteins used in this study.** Proteins were heterologously expressed in *E. coli* and purified using amylose affinity chromatography. $FusC_{op}$ means the FusC protein was derived from an *E. coli*-optimized FusC gene.[5] MBP (43 kDa) is observed as truncated product in FusB and $FusC_{op}$.
